# A proteogenomic surfaceome study identifies DLK1 as an immunotherapeutic target in neuroblastoma

**DOI:** 10.1101/2023.12.06.570390

**Authors:** Amber K. Weiner, Alexander B. Radaoui, Matthew Tsang, Daniel Martinez, Simone Sidoli, Karina L. Conkrite, Alberto Delaidelli, Apexa Modi, Jo Lynne Rokita, Khushbu Patel, Maria V. Lane, Bo Zhang, Chuwei Zhong, Brian Ennis, Daniel P. Miller, Miguel A. Brown, Komal S. Rathi, Pichai Raman, Jennifer Pogoriler, Tricia Bhatti, Bruce Pawel, Tina Glisovic-Aplenc, Beverly Teicher, Stephen W. Erickson, Eric J. Earley, Kristopher R. Bosse, Poul H. Sorensen, Kateryna Krytska, Yael P. Mosse, Karin E. Havenith, Francesca Zammarchi, Patrick H. van Berkel, Malcolm A. Smith, Benjamin A. Garcia, John M. Maris, Sharon J. Diskin

## Abstract

Cancer immunotherapies have produced remarkable results in B-cell malignancies; however, optimal cell surface targets for many solid cancers remain elusive. Here, we present an integrative proteomic, transcriptomic, and epigenomic analysis of tumor specimens along with normal tissues to identify biologically relevant cell surface proteins that can serve as immunotherapeutic targets for neuroblastoma, an often-fatal childhood cancer of the developing nervous system. We apply this approach to human-derived cell lines (N=9) and cell/patient-derived xenograft (N=12) models of neuroblastoma. Plasma membrane-enriched mass spectrometry identified 1,461 cell surface proteins in cell lines and 1,401 in xenograft models, respectively. Additional proteogenomic analyses revealed 60 high-confidence candidate immunotherapeutic targets and we prioritized Delta-like canonical notch ligand 1 (DLK1) for further study. High expression of DLK1 directly correlated with the presence of a super-enhancer spanning the DLK1 locus. Robust cell surface expression of DLK1 was validated by immunofluorescence, flow cytometry, and immunohistochemistry. Short hairpin RNA mediated silencing of DLK1 in neuroblastoma cells resulted in increased cellular differentiation. ADCT-701, a DLK1-targeting antibody-drug conjugate (ADC), showed potent and specific cytotoxicity in DLK1-expressing neuroblastoma xenograft models. Moreover, DLK1 is highly expressed in several adult cancer types, including adrenocortical carcinoma (ACC), pheochromocytoma/paraganglioma (PCPG), hepatoblastoma, and small cell lung cancer (SCLC), suggesting potential clinical benefit beyond neuroblastoma. Taken together, our study demonstrates the utility of comprehensive cancer surfaceome characterization and credentials DLK1 as an immunotherapeutic target.

**Highlights:**

- Plasma membrane enriched proteomics defines surfaceome of neuroblastoma
- Multi-omic data integration prioritizes DLK1 as a candidate immunotherapeutic target in neuroblastoma and other cancers
- DLK1 expression is driven by a super-enhancer
- *DLK1* silencing in neuroblastoma cells results in cellular differentiation
- ADCT-701, a DLK1-targeting antibody-drug conjugate, shows potent and specific cytotoxicity in DLK1-expressing neuroblastoma preclinical models

## Introduction

Neuroblastoma remains one of the deadliest cancers in children. Despite intense multimodal therapy, the survival rate for high-risk neuroblastoma is less than 50% and relapse is generally incurable ^1,2^. Sequencing of neuroblastoma tumors obtained at diagnosis has revealed a relative paucity of recurrent somatic driver mutations ^3–6^. Approximately 15% of high-risk neuroblastomas harbor an activating mutation in the Anaplastic Lymphoma Kinase (*ALK*) gene ^7–10^, and the presence of an *ALK* aberration is associated with poor survival probability ^11^. Lorlatinib, a 3^rd^ generation allosteric ALK inhibitor has shown promising anti-tumor activity in patients with relapsed disease harboring an *ALK* mutation ^12^ and is now being evaluated in newly diagnosed patients (ClinicalTrials.gov: NCT03126916). However, beyond *ALK*, no other kinase has been amenable to enzymatic inhibition in high-risk neuroblastoma. Monoclonal antibody-based immunotherapy targeting GD2, a disialoganglioside on the cell surface of neuroblastoma, extended survival ^13–15^, leading to the only licensed immunotherapy for a pediatric solid malignancy. However, anti-GD2 therapy is associated with severe side effects due to GD2 expression on pain fibers and patients often relapse during or after treatment ^13,16^. Several other surface proteins have been nominated as candidate immunotherapeutic targets in neuroblastoma, including L1CAM ^17^, ALK ^7,18–20^, GPC2 ^21^, DLL3 ^22^ and B7H3 ^23,24^; however, no non-GD2 directed immunotherapy has to date shown significant anti-tumor activity in patients ^25^.

Despite recent advances in cancer immunotherapy that have enabled targeted drug delivery to tumor cells through internalization of cytotoxic payloads ^26^, the identification of optimal cancer cell surface proteins with low or absent expression on normal tissues remains a challenge. ADCs and chimeric antigen receptor (CAR) T-cell immunotherapy require identification of suitable cell surface epitopes for effective binding to occur. Experimental follow up on multiple candidate targets is needed since prioritized proteins may fail at any point during validation and pre-clinical studies. In addition, identification of proteins that can be targeted in combination would help overcome single agent-associated liabilities such as antigen loss and tumor heterogeneity. Previous studies to discover novel immunotherapeutic targets in childhood cancers have relied heavily on mining mRNA expression data for identification ^21,27–32^. However, RNA-based strategies do not account for the rate of translation, RNA turnover, and localization of the final protein product, which can result in potentially missed targets ^33–35^. Recent instrumentation and computational advances have enabled mass spectrometry-based proteomics to emerge as a reliable method to identify and quantify thousands of proteins simultaneously in an unbiased fashion, ranging from subcellular fractions to full proteomes ^36–38^. Here, we present an integrative proteomic, transcriptomic and epigenomic approach to prioritize candidate cell surface proteins to serve as immunotherapeutic targets in neuroblastoma. This is followed by extensive validation and pre-clinical studies to credential DLK1, a prioritized surface protein, as an immunotherapeutic target in neuroblastoma.

## Results

### Mass spectrometry-based surfaceome of neuroblastoma

To quantify the repertoire of cell surface proteins (surfaceome) on neuroblastomas, we first optimized a sucrose gradient plasma membrane enrichment methodology previously used in other cancers ^39^. Enriched plasma membrane proteins were then subjected to nano-liquid chromatography coupled to mass spectrometry (nLC-MS/MS). We applied this approach to a genetically diverse panel of human-derived neuroblastoma cell lines (N=9) and xenograft (N=12) tumor models **(Table 1)**. A total of 7,172 proteins in cell lines and 6,831 proteins in xenograft models were detected and quantified by mass spectrometry. To obtain a high-confidence set of surface proteins, quantified proteins were filtered using COMPARTMENTS (Compartments > 3) ^40^ and the surface protein consensus score (SPC) from CIRFESS (SPC > 0) ^41^, yielding 1,461 surface proteins in cell lines and 1,401 in xenograft models, respectively. Proteins that did not pass these strict thresholds were evaluated using GOrilla ^42^ to determine which cell compartments were enriched. The top GO term for filtered out proteins was extracellular region and extracellular space (Cell Line_ExtracellularRegion_: p=1.2×10^−10^; q=1.9×10^−7^; Xenograft_ExtracellularSpace_: p=2.63×10^−11^; q=4.16×10^−8^), thus excluding secreted proteins. To assess reproducibility, quantified surface proteins were compared across biological and technical replicates. We observed strong reproducibility between biological and technical replicates for cell lines and xenografts (Cell Line: protein overlap range: 82%-90%; Pearson’s R: 0.88-0.96; Xenograft: protein overlap range: 67%-86%; Pearson’s R: 0.89-0.96) (**Figures 1A and 1B; Figures S1A and S1B; Table 1)**. We then queried this mass-spectrometry-based protein list for surface proteins being developed, or previously studied, as candidate immunotherapeutic targets in neuroblastoma ^21,22,43–46^. We observed NCAM1, L1CAM, GPC2, CD276, SLC6A2, DLL3 and ALK among the most abundant proteins in both cell lines and xenograft models, validating our approach **(Figures 1C-1F).**

**Figure 1.**
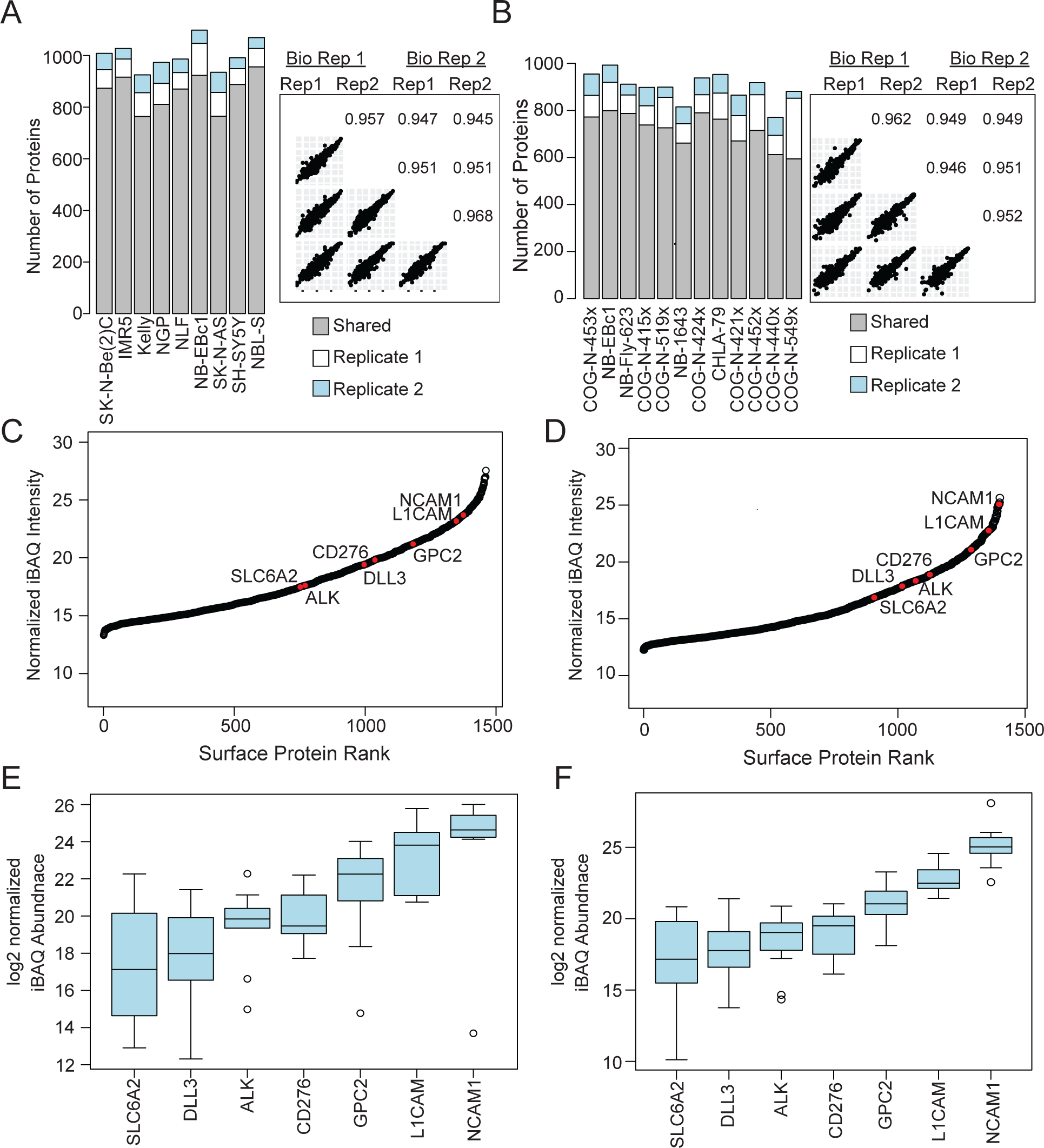
Sucrose density gradient methodology coupled to mass spectrometry defines the neuroblastoma surfaceome. **(A-B)** Bar plots show the reproducibility of protein identification between two biological replicates for cell lines **(A)** and xenograft models **(B)**. Correlation between biological and technical replicates of the same model are calculated. **(C-D)** Proteins are ranked based on mean normalized iBAQ abundance with the most abundant proteins on the right of the swoosh plot. Known cell surface proteins in development as immunotherapeutic targets (highlighted in red) are among the most abundant proteins detected by mass spectrometry in neuroblastoma cell lines **(C)** and xenograft models **(D)**. **(E-F)** Box and whisker plots illustrating the abundance of known surface proteins highlighted in red in panels C-D across cell lines and xenografts. See related **Figure S1.**

**Table 1.** Models for mass spectrometry surfaceome analysis. Overview of samples analyzed by mass spectrometry. Each sample’s *MYCN* status (amplified or not amplified), *ALK* status (wild type or mutated), phase of therapy (Diagnosis, Progressing Disease or Progressive Disease – Post-Mortem PD-PM), and model type are annotated. Average correlation between technical and biological replicates, RNA-protein correlation and number of replicates are indicated. PDX: patient-derived xenograft; CDX: cell line derived xenograft.

| Model Name | Model Type | *MYCN* Status | *ALK* Status | Phase of Therapy | MS Replicate Correlation | RNA-Protein Correlation | Number of Biological Replicates |
| --- | --- | --- | --- | --- | --- | --- | --- |
| IMR-05 | Cell Line | Amplified | WT | Diagnosis | 0.908 | 0.54 | 2 |
| Kelly | Cell Line | Amplified | F1174L | Unknown | 0.944 | 0.54 | 2 |
| NGP | Cell Line | Amplified | WT | Unknown | 0.927 | 0.54 | 2 |
| NLF | Cell Line | Amplified | WT | Diagnosis | 0.947 | 0.48 | 3 |
| SK-N-Be(2)C | Cell Line | Amplified | WT | Progressing | 0.934 | 0.62 | 2 |
| NB-Ebc1 | Cell Line | Not Amplified | WT | Progressing | 0.879 | 0.52 | 2 |
| NBL-S | Cell Line | Not Amplified | WT | Diagnosis | 0.957 | 0.53 | 2 |
| SH-SY5Y | Cell Line | Not Amplified | F1174L | Progressing | 0.950 | 0.52 | 3 |
| SK-N-AS | Cell Line | Not Amplified | WT | Progressing | 0.890 | 0.48 | 3 |
| COG-N-415x | PDX | Amplified | F1174L | Progressing | 0.963 | 0.52 | 3 |
| COG-N-421x | PDX | Amplified | WT | PD-PM | 0.924 | 0.42 | 2 |
| COG-N-424x | PDX | Amplified | WT | Diagnosis | 0.942 | 0.47 | 2 |
| COG-N-440x | PDX | Amplified | WT | PD-PM | 0.929 | 0.43 | 3 |
| COG-N-452x | PDX | Amplified | F1174L | PD-PM | 0.891 | 0.43 | 4 |
| COG-N-453x | PDX | Amplified | F1174L | PD-PM | 0.929 | 0.44 | 2 |
| COG-N-519x | PDX | Amplified | WT | PD-PM | 0.920 | 0.53 | 4 |
| NB-1643 | CDX | Amplified | R1275Q | Diagnosis | 0.938 | 0.47 | 2 |
| CHLA-79 | CDX | Not Amplified | WT | Progressing | 0.929 | 0.48 | 2 |
| COG-N-549x | PDX | Not Amplified | F1174L | Progressing | 0.912 | 0.50 | 4 |
| NB-EBc1 | CDX | Not Amplified | WT | Progressing | 0.929 | 0.49 | 2 |
| NB-Fly-623 | PDX | Not Amplified | R1275Q | PD-PM | 0.951 | 0.46 | 2 |

It has been observed that RNA and protein are discordant due to many processes including RNA and protein modifications, molecule stability, rate of translation and protein degradation ^47^. To assess the correlation between RNA and protein for the quantified surface proteins, we utilized matched RNA-sequencing data for both cell lines and xenograft models ^48,49^. Of the evaluable proteins, 63% (771/1,220) of surface proteins in cell lines and 43% (489/1165) in xenograft models showed a positive correlation (Pearson’s R Correlation > 0.4). (**Figures S1C and S1D**). This observation for surface proteins in neuroblastoma is consistent with the mass spectrometry-based draft of the human proteome and recent literature ^50,51^. Notably, this approach allowed us to identify both positively and negatively correlated proteins (**Figure S1E**).

To identify highly expressed proteins detected at low abundance by RNA-sequencing, we first examined the distribution of mass spectrometry (normalized iBAQ) and RNA-sequencing (log2(TPM)) post filtering and found that mass spectrometry data was normally distributed while RNA was skewed to the left due to low level RNAs. Therefore, we applied a gaussian mixture model and identified two distributions in the RNA-sequencing data for each cell line and xenograft model (**Figure S1F**). We extracted transcripts that were below the mean of the lower distribution and cross referenced them with the mass spectrometry data. This process identified recurrent proteins that were low by RNA using a Gaussian Mixture Model yet identified at the protein level (**Figures S1G and S1H**).

### Multi-omic data integration prioritizes surfaceome proteins as candidate immunotherapeutic targets

To identify the most promising cell surface proteins for further validation, we developed a multi-omic data integration and prioritization pipeline and applied it independently to cell lines and xenograft models (**Figures 2A and 2B**). After applying the COMPARTMENTS and SPC criteria to restrict to surface proteins, 1,461 proteins remained in cell lines and 1,401 proteins in xenograft models, respectively. We then restricted to proteins with low mRNA expression in normal tissues compared to neuroblastoma as measured by RNA-sequencing data from Genotype-Tissue Expression (GTEx) ^52,53^ and excluded those with a fold change (FC) of neuroblastoma compared to each normal tissue <4 in four or more tissues excluding brain and reproductive organs (N_CellLine_ = 177, N_Xenograft_ = 199; see **Methods**). Next, proteins were evaluated for MS abundance, retaining proteins with abundance >50^th^ percentile when averaged across all cell lines or xenograft models, respectively (N_CellLine_ = 45; N_Xenograft_ = 55). This resulted in 60 unique proteins with high expression in neuroblastoma cell lines and/or xenograft models and minimal or no expression in selected normal tissues.

**Figure 2.**
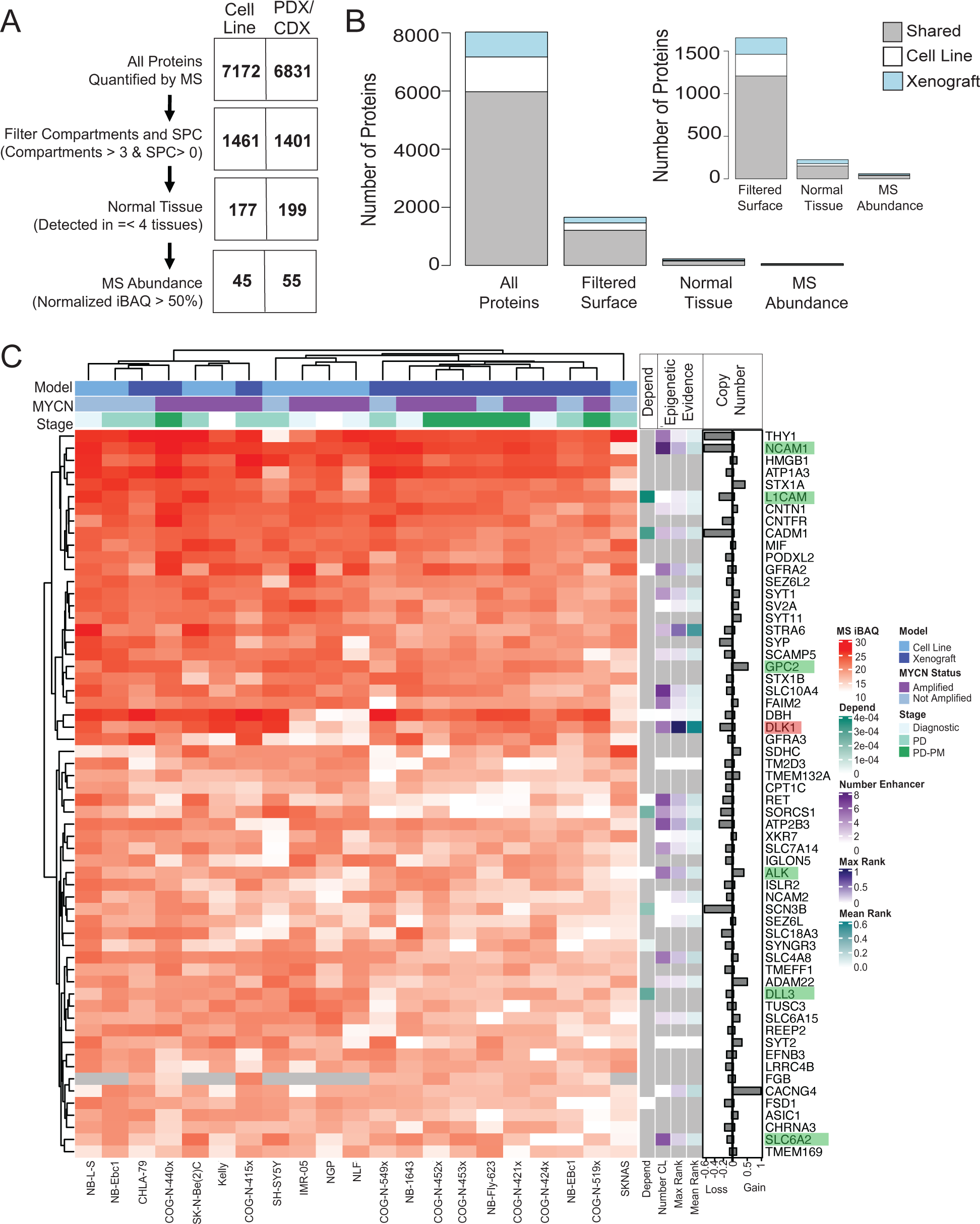
Proteogenomic data integration prioritizes potential immunotherapeutic targets. **(A)** Schematic showing integration of mass spectrometry and RNA-sequencing to prioritize cell surface proteins as candidate immunotherapeutic targets. Proteins were filtered using the COMPARTMENTS and SPC annotation, normal tissue expression and MS abundance. High concordance was observed between neuroblastoma cell line and PDX models **(B)** Number of proteins identified in both cell lines and xenografts individually and shared in both for each step of prioritization **(C)** Annotated heatmap of top-prioritized surface proteins across neuroblastoma cell lines and xenograft models. The heatmap displays the normalized MS iBAQ intensity. For each sample, model (cell line or xenograft), MYCN status and phase of therapy are indicated above heatmap. Dependency status, epigenetic evidence, and DNA copy number are summarized to the right of the heatmap. Known targets are highlighted in green and our prioritized target DLK1 is highlighted in red.

We sought to further characterize proteins and determine a potential underlying genomic or epigenetic mechanism driving their expression. Finding no recurrent mutations associated with over-expression of the more abundant proteins recovered, we focused on epigenetic mechanisms. Recent studies have shown that core regulatory circuitry (CRC) transcription factors bind at super enhancers to control gene expression and maintain cell state in neuroblastoma ^54–58^. Therefore, we first evaluated H3K27ac ChIP-sequencing data on a panel of ten neuroblastoma cell lines ^59^. Overall, a putative super enhancer was identified for 3,206 genes across the ten cell lines. The number of cell lines and intensity of the H3K27Ac were overlaid with the mass spectrometry data of cell lines and xenografts for the top 60 targets that passed our normal tissue threshold. Prioritized proteins were visualized and overlaid with genomic and epigenomics data (**Figure 2C**).

### DLK1 is a cell surface protein with expression driven by a super enhancer

We selected the Delta Like Non-Canonical Notch Ligand 1 gene, *DLK1*, as a novel top-prioritized protein in our multi-omic surfaceome due to multiple levels of support. A high correlation between RNA and protein expression was observed in neuroblastoma cell lines (R=0.97, p=1.1×10^−3^) with a modest correlation in xenograft models (r=0.42, p=0.18) (**Figures 3A and 3B)**. Examination of RNA-sequencing data from neuroblastoma patient tumors and diverse normal tissues from GTEx showed *DLK1* mRNA expression to be absent in the majority of normal tissues, except the adrenal, pituitary glands and ovary (**Figure 3C**). *DLK1* expression was associated with *MYCN* amplification status in neuroblastoma (TARGET high-risk: ANOVA p=6.45×10^−3^, Kocak Stage 4: ANOVA p=0.024, SEQC high-risk: ANOVA p=0.014) (**Figure S2A**). Further interrogation of matched whole genome sequencing (WGS) data for high-risk patient tumors revealed no focal amplifications involving *DLK1*, suggesting that the major mechanism of DLK1 overexpression in a large subset of neuroblastomas is epigenetic.

**Figure 3.**
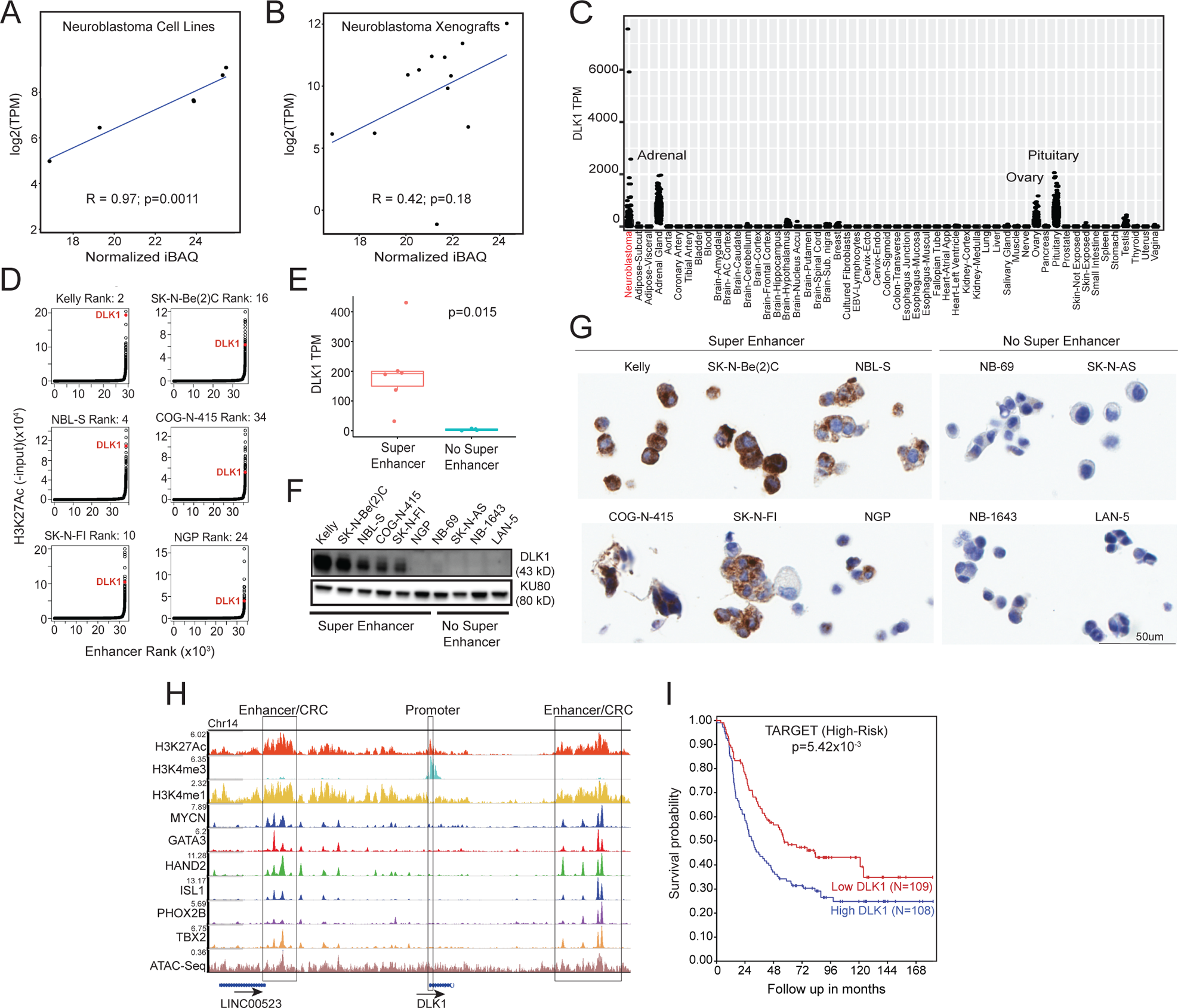
Prioritization of DLK1 as a candidate immunotherapeutic target in neuroblastoma. **(A-B)** DLK1 expression correlation in cell line and xenograft models comparing mass spectrometry and RNA sequencing. **(C)** DLK1 RNA-sequencing data indicate high expression in a subset of neuroblastoma and normal tissue expression restricted to the adrenal/pituitary gland and ovary. **(D)** Swoosh plot of H2K27ac ChIP-sequencing signal show a subset of neuroblastoma cell lines have a super enhancer as indicated in red. **(E-G)** Active super-enhancer correlates with higher DLK1 expression measured by RNA-sequencing, immunoblotting and immunohistochemistry of a neuroblastoma cell plug array. **(H)** Visualization of DLK1 locus highlighting H3K27Ac, H3K4me1 and H3K4me3 and CRC transcription factors in SK-N-Be(2)C. **(I)** High DLK1 mRNA expression is associated with poor overall survival in high-risk neuroblastoma. See related **Figure S2**.

Analysis of histone ChIP-sequencing for H3K27ac revealed the presence of a super-enhancer at the *DLK1* locus in six of the ten neuroblastoma cell lines evaluated (**Figures 3D** and **S2B**). Indeed, the *DLK1* locus ranked as one of the strongest super-enhancers across the panel of cell lines (rank 2 to 34; percentage rank: >0.99; **Figures S2C and S2D**). To investigate if this super-enhancer is associated with high expression of DLK1, we first compared RNA-expression levels of cell lines with and without the super enhancer. Cell lines harboring the active super-enhancer had significantly higher levels of *DLK1* mRNA (p = 0.015; **Figure 3E**). These results were subsequently confirmed at the protein level using immunoblotting (**Figure 3F**) and immunohistochemistry of neuroblastoma cells (**Figure 3G**). Finally, we visualized ChIP-sequencing data for the CRC transcription factors, including MYCN, GATA3, HAND2, ISL1, PHOX2B and TBX2. These CRC transcription factors were observed to bind up and downstream of the *DLK1* gene locus (**Figure 3H**), suggesting *DLK1* may be regulated by the CRC in neuroblastoma.

### *DLK1* is expressed in patient samples and high levels associate with poor survival

We next examined whether high levels of *DLK1* mRNA correlate with high-risk neuroblastoma outcome. Analysis of RNA-sequencing data restricted to high-risk neuroblastoma patient tumors sequenced through the TARGET project showed that high *DLK1* expression is associated with a worse overall survival p= 5.42×10^−3^; **Figure 3I**). In two independent datasets, high expression trended with poorer outcome but did not reach statistical significance (SEQC p=0.071; Kocak p = 0.057; **Figure S2E**).

### DLK1 surface expression validates by immunofluorescence, flow cytometry, and immunohistochemistry

To assess whether DLK1 is amenable for immunotherapeutic targeting, we sought to validate its expression on the surface of neuroblastoma cells. We first performed immunofluorescence (IF) on a panel of four genetically diverse neuroblastoma cell lines with either high endogenous protein expression of DLK1 (SK-N-Be(2)C and Kelly) or low/no DLK1 expression (NB-69 and NLF). We observed that DLK1 co-localized with cadherin, a known cell surface protein, in DLK1 positive cells but not in cells lacking DLK1 expression (**Figure 4A**). In parallel, flow cytometry in the same panel of neuroblastoma cell lines further confirmed cell surface expression of DLK1 is restricted to cell lines with high *DLK1* mRNA expression (**Figure 4B**).

**Figure 4.**
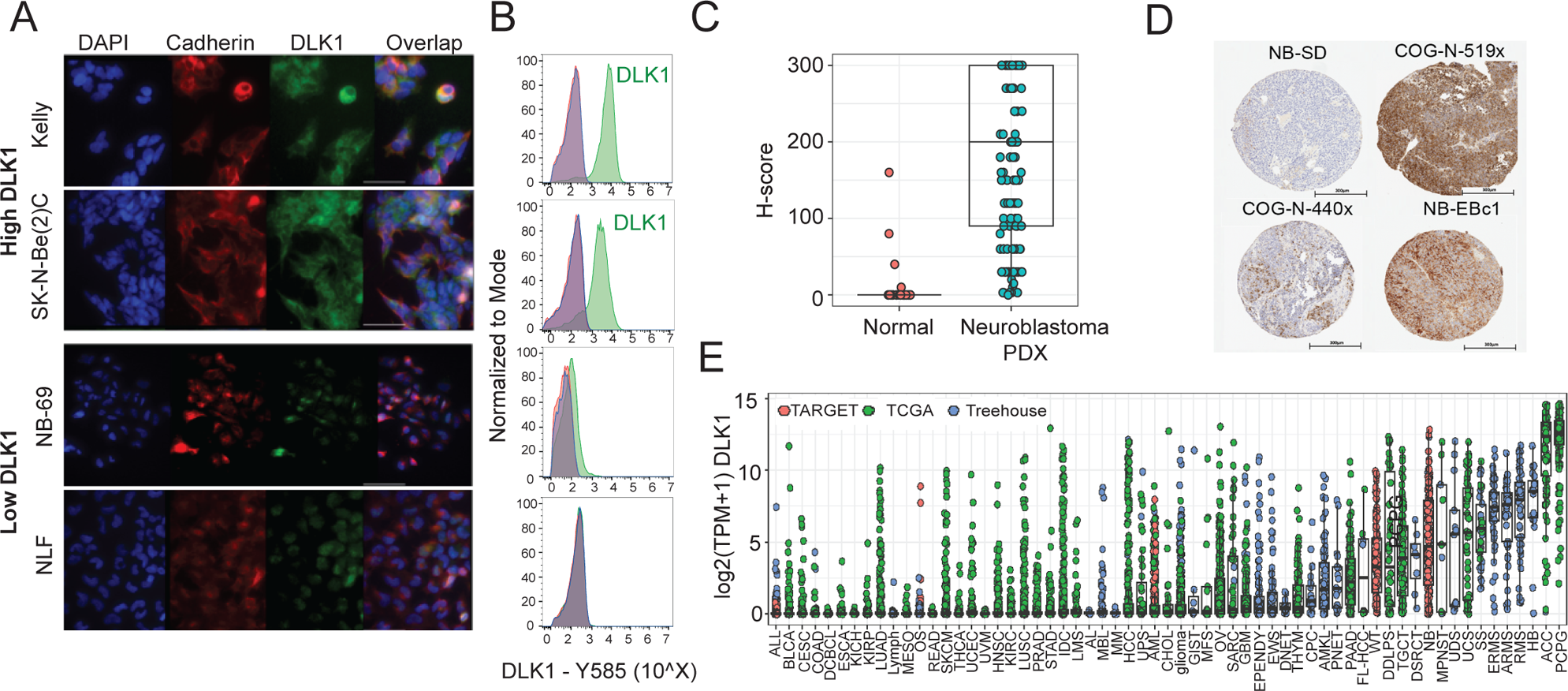
Validation of DLK1 as a cell surface protein differentially expressed in neuroblastoma compared to normal tissues. **(A)** Immunofluorescence shows colocalization of DLK1 (green) with the cell surface marker Cadherin (red) in cell lines with high DLK expression. **(B)** Flow cytometry validates cell surface expression in a panel of neuroblastoma cell lines positive for DLK1 and negative (no endogenous expression) **(C)** Immunohistochemistry using tissue microarrays of xenograft neuroblastoma and pediatric normal tissues validates preferential expression of DLK1 in neuroblastoma. H-score shows differential expression between neuroblastoma and normal tissues. **(D)** Representative neuroblastoma cores with high and low DLK1 expression. **(E)** Expression of DLK1 in Treehouse Childhood Cancer Initiative dataset. See related **Figure S3**.

We next performed additional immunohistochemistry (IHC) staining on both a pediatric normal tissue array (N=41 tissues) and a neuroblastoma xenograft (cell and patient) array (N=32 tumors in duplicate). To evaluate expression and heterogeneity, we employed a modified H-score, calculated by multiplying the staining intensity times the percent of stained cells resulting in a scale ranging from 0-300. Overall, 81.6% of xenograft tumors had a H-score above 50. Significant differential protein expression of DLK1 was observed between normal pediatric tissues and neuroblastoma (**Figures 4C and S3A)**, with DLK1 positive and negative xenograft examples visualized **(Figure 4D**). DLK1 expression was seen in adrenal (1 month age patient) and pituitary cores (70 months age patient); (**Figure S3B)**. We also performed IHC on a panel of neuroblastoma tumors. While the antibody did not perform as robustly in archival tumor tissues as xenograft samples (Xenograft_Median_ = 200; Tumor_Median_ = 10), neuroblastomas showed significantly higher expression than most normal tissues, excepting the adrenal gland. (**Figure S3C)**. Other normal tissues showed negligible staining. Next, we performed additional staining on the whole adrenal glands to determine the localization of DLK1 expression, which showed strong staining (H-Score: 200-300) in the medullary chromaffin cells, but no staining in the adrenal cortex (**Figures S3C and S3D**).

To identify other histologies that could potentially benefit from a DLK1-directed therapy, we utilized RNA-sequencing data from The Cancer Genome Atlas (TCGA) and Treehouse Childhood Cancer Initiative to examine DLK1 expression across a panel of adult and pediatric solid tumors. Pheochromocytoma/paraganglioma, adrenocortical carcinoma, hepatoblastoma and several sarcomas had the highest expression of DLK1 along with neuroblastoma, with several other histotypes demonstrating high expression in a subset of samples (**Figure 4E**).

### *DLK1* depletion promotes neuroblastoma cellular differentiation

Based on prior reports ^60–62^, we sought to further explore the role of high DLK1 expression in neuroblastoma cellular differentiation. We first treated neuroblastoma cell lines SK-N-Be(2)C (*MYCN* Amplified) and SH-SY5Y (*MYCN* not amplified) with all-trans retinoic acid (ATRA), which is known to cause these cells to terminally differentiate ^63,64^. We observed decreased DLK1 protein expression in response to ATRA-induced differentiation in both cell models (**Figures 5A and 5B**). Next, we performed shRNA-mediated silencing of *DLK1* in SK-N-Be(2)C cells, which express high endogenous levels of *DLK1*. Depletion of *DLK1* using two distinct shRNAs targeting *DLK1* was confirmed by Western blot and showed a 95% decrease in DLK1 expression compared to shScrambled control in SK-N-Be(2)C cells (**Figures 5C, and S4A**). Following transduction, we observed neurite formation in the sh*DLK1* treated cells but not the control cells (**Figure 5C**). To further quantify this phenotype, we assessed neurite length and cell growth as a measure for differentiation and proliferation, respectively. SK-N-Be(2)C cells were plated in 96-well format and transductions were performed in two biological and six technical replicates for scrambled control and *DLK1*-directed shRNA silencing. We observed significantly increased neurite length relative to cell body in the sh*DLK1* cells compared to controls (p = 3.16×10^−6^; **Figures 5D, S4B, and S4C**) with no differences observed in cell growth. Finally, we performed a full proteome analysis on SK-N-Be(2)C shScrambled (control) and sh*DLK1* treated cells in duplicate. A total of 5,240 proteins were detected and quantified. DLK1 was the fourth most down-regulated protein when comparing sh*DLK1* treated cells to shScrambled control cells (log_2_ fold change: −3.86). Ingenuity pathway analysis (IPA) of differentially expressed proteins (n=976) confirmed enrichment in processes involving neuron and neurite formation (p = 3.37×10^−3^ to 8.23×10^−3^; **Figure 5E**). Taken together, these data support the literature that DLK1 expression plays an important role in suppressing cellular differentiation in neuroblastoma and likely maintaining an adrenergic cellular state ^54–58^.

**Figure 5.**
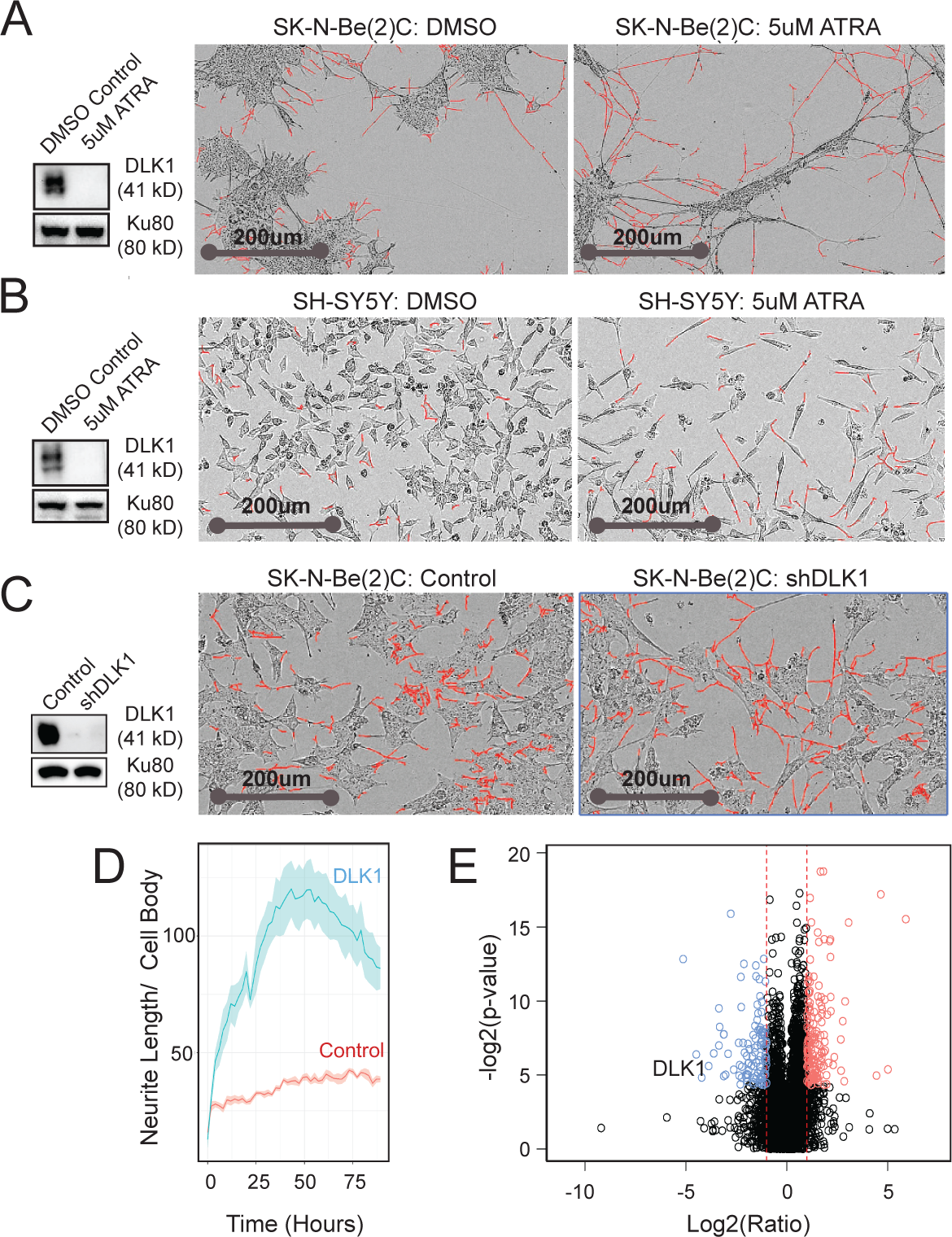
DLK1 is a biologically relevant protein that plays a critical role in cellular differentiation. **(A-B)** SK-N-Be(2)C and SH-SY5Y neuroblastoma cells treated with all trans retinoid acid (ATRA) exhibited depleted expression of DLK1 and induced differentiation. Images of SK-N-Be(2)C and SH-SY5Y cells treated with DMSO alone and 5uM ATRA **(C)** Western blotting demonstrating robust knockdown of DLK1 compared to scrambled control. Images of scrambled control and DLK1 knockdown in SK-N-Be(2)C cells **(D)** Incucyte NeuroTrack analysis shows differentiation phenotype is associated with an increase in neurite length following shRNA DLK1 knockdown **(E)** Mass spectrometry analysis of Control and shDLK1 cells indicates DLK1 is the fourth most downregulated protein in the proteome. See related **Figure S4**.

### ADCT-701, a DLK1-targeted ADC, shows potent and specific antitumor activity in human neuroblastoma xenograft models

To therapeutically target DLK1, we utilized two approaches, a monoclonal antibody targeting DLK1 with and without a pyrrolobenzodiazepine (PBD) dimer as payload. We first studied the DLK1-targeting monoclonal antibody HuBA-1-3D, a humanized IgG1 antibody directed against human DLK1 (not mouse cross-reactive), for antitumor efficacy in four neuroblastoma xenograft models with high DLK1 expression but saw no evidence of anti-tumor efficacy (**Figure S5A)**. Next, we tested ADCT-701, an ADC composed of HuBA-1-3D conjugated using GlycoConnect^TM^ technology to PL1601, which contains HydraSpace^TM^, a Val-Ala cleavable linker and the potent PBD dimer cytotoxin SG3199 ^65^ **(Figure 6A)**. Prior to efficacy testing, we confirmed that a single dose of ADCT-701 at 1 mg/kg caused no obvious toxicity in mice including no weight loss. Moreover, pharmacokinetic (PK) analysis of a single dose at 1 mg/kg in non-tumor bearing mice showed that the profiles of total and conjugated antibody (drug to-antibody ratio [DAR] ≥1.9) were comparable, indicating high *in vivo* stability ^65^.

**Figure 6.**
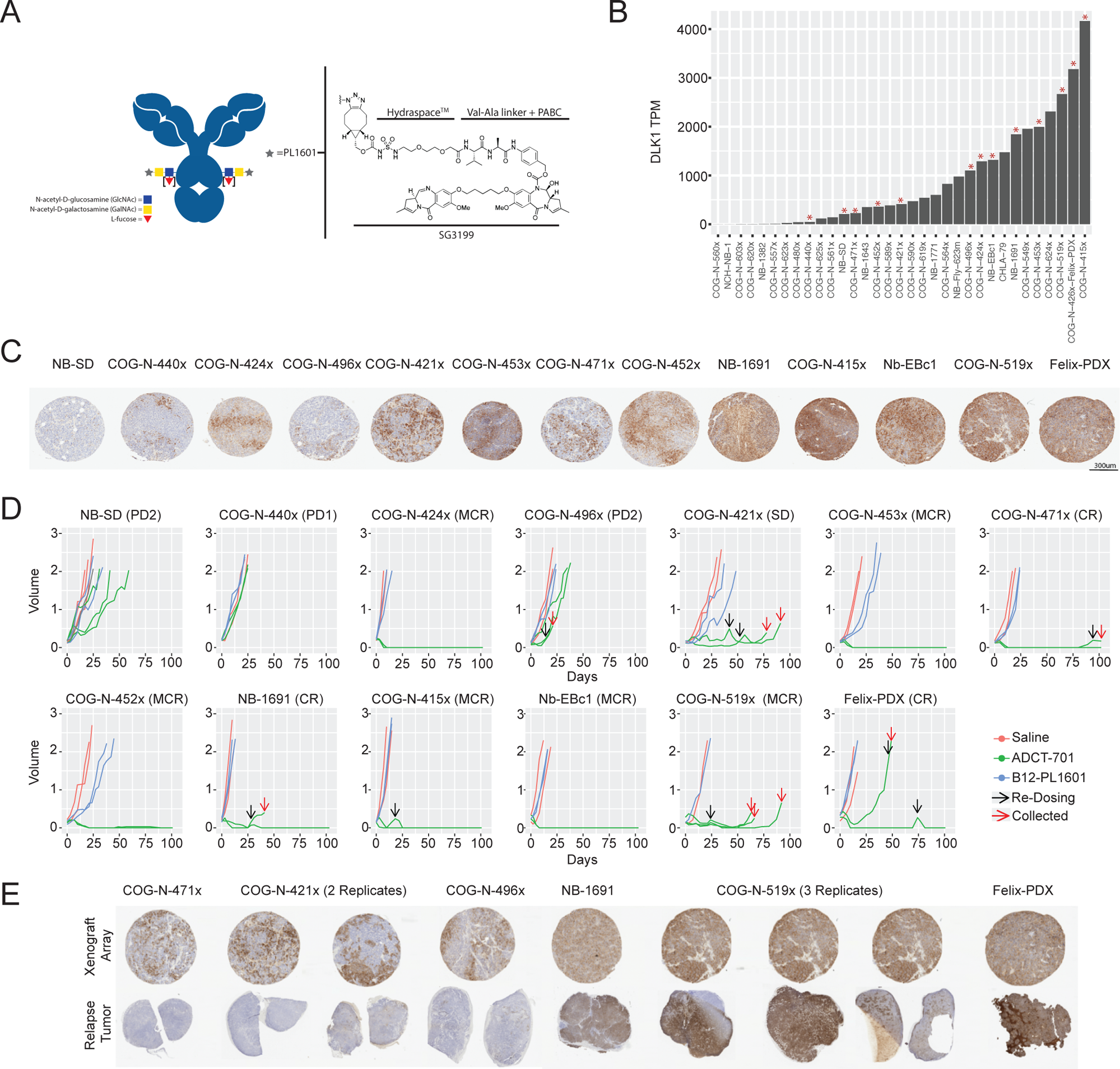
ADCT-701 shows specific and potent anti-tumor activity in neuroblastoma models expressing DLK1. **(A)** ADCT-701 drug structure. **(B)** DLK1 RNA expression across neuroblastoma xenograft models. Red asterisks indicate models used for ADCT-701 efficacy studies **(C)** DLK1 IHC staining of a xenograft array show variable DLK1 expression across models **(D)** Tumor volumes (cm3) were measured for the saline control (red), non-targeting control ADC – B12-PL1601 (blue) and ADCT-701 (green). Arrows indicate the administration of a second dose (black) and tumor collection (red). Each model was scored based on PIVOT guidelines (PD1/PD2-Progressive Disease; SD-Stable Disease; CR-Complete Response; MCR-Maintained Complete Response) **(E)** IHC of CDX/PDX array (top row) are compared to the relapse tumors (bottom row). See related **Figure S5**.

To test the efficacy of ADCT-701, we selected thirteen xenograft tumor models based on high-, moderate- and low-DLK1 expression as determined by IHC and RNA-sequencing (**Figures 6B**, 6C**, and S5B**). Mice harboring established xenografts (tumor volume ∼ 200 mm^3^) were randomized to receive a single 1mg/kg dose of ADCT-701, B12-PL1601 (a non-binding isotype-control antibody targeting HIV gp120 conjugated to the PL1601 payload following the same procedure described for ADCT-701) or saline administered by tail vein, with two or three mice per arm using standard methodology utilized in the National Cancer Institute Pediatric Preclinical In Vivo Testing (PIVOT) Program ^66^. High expressing models including Felix-PDX, COG-N-519x, Nb-EBC1, COG-N-415x, NB-1691 and COG-N-452x exhibited complete responses, many of which were maintained throughout the observation period of 100 days. NB-SD and COG-N-440x have low expression of DLK1 and showed progressive disease. Therefore, ADCT-701 showed potent and specific anti-tumor activity that correlated with DLK1 surface expression in preclinical models of neuroblastoma (**Figures 6D and S5C**). No toxicities or weight loss were observed. PK analysis of ADCT-701 in non-tumor-bearing mice following a single intravenous administration of ADCT-701 at 1 mg/kg showed the profiles of total antibody and ADC (DAR ≥1) were comparable, indicating high *in vivo* stability (**Figure S5D**).

Six of the thirteen models showed tumor regrowth in at least one mouse after regressions induced by ADCT-701 (**Figure 6E**). Two tumors (one COG-N-415x and one Felix-PDX) showed a second complete remission following a second dose of ADCT-701, suggesting that the tumors retained DLK1 expression (**Figure 6D**). Initially COG-N-421x showed stable disease and COG-N-471x a complete response despite heterogeneous expression at study entry. This can be explained by the bystander effect, which can occur when an ADC is internalized by a target-expressing cell and has its payload released intracellularly with subsequent diffusion of the payload out of the target cell to adjacent cells where it can induce cytotoxicity independent of target antigen expression ^67^. To determine DLK1 expression in ADC-relapsed tumors, they were harvested before they reached tumor burden endpoint. These relapsed tumors showed either transient response or stable disease when retreated with ADCT-701, and harvested tumors following a second ADC treatment showed no DLK1 expression by IHC (COG-N-471x, COG-N-421x, COG-N-496x). For models with higher DLK1 expression that initially responded to ADCT-701 treatment, tumors that regrew after a second dose showed robust DLK1 staining. COG-N-519x and NB-1691 continued to grow after second dose and were DLK1-positive by IHC, suggesting resistance to the drug. COG-N-519x is p53 mutated and NB-1691 has a MDM2 amplification that correlate with PBD sensitivity ^21^. More detailed statistical analyses can be found in **Figure S5C**. Taken together, ADCT-701 showed potent and specific activity in aggressive models of neuroblastoma.

## Discussion

In this study, we performed an unbiased survey of cell surface proteins (surfaceome) in neuroblastoma to facilitate the identification of candidate immunotherapeutic targets. Sucrose gradient ultracentrifugation was utilized for sub-cellular fractionation and protein extraction from cell line and xenograft models. Through plasma membrane protein enrichment coupled to mass spectrometry-based proteomics, surface proteins were quantified directly rather than relying on RNA-based inferences of plasma membrane expression. To identify optimal surface proteins suitable for immunotherapy development, we employed a multi-omic approach for target prioritization. Specifically, we integrated cancer and normal tissue expression (RNA-sequencing and MS-based proteomics), investigated genomic and epigenetic mechanisms of expression (WGS and ChIP-sequencing) and evaluated potential neuroblastoma dependency through DepMap (RNAi/CRISPR-screening). To assess mechanisms of over-expression, DNA copy number gain and super-enhancer status were evaluated. Assessing epigenetic processes including the acquisition of a super enhancer provided an important understanding of potential drivers of high *DLK1* expression, which differs from previously identified immunotherapeutic targets. Overall, this approach was able to prioritize known immunotherapeutic targets ^21,28^ and identify additional cell surface proteins for validation and development, including DLK1.

DLK1 is a cell surface protein that is thought to suppress Notch signaling, which regulates critical cellular processes including proliferation, development, and differentiation ^62,68^. The *DLK1* gene is located at an imprinted region within chromosome band 14q32 and expression is derived from the paternal allele in normal fetal development ^69^. In the current study, tissue microarray (TMA) staining of tumor and normal tissues for DLK1 showed normal tissue expression restricted to the adrenal medulla and pituitary gland. In neuroblastoma, DLK1 may contribute to the undifferentiated state ^61,70^, similar to other cell types and cancers reviewed by Pittaway and colleagues ^71^. More specifically, depletion of DLK1 was associated with differentiation, while over expression inhibited this process ^61,62,70^. Our study added to these observations by using the Incucyte zoom to assess cell growth and neurite length following shRNA knockdown or ATRA treatment. We further identified a previously unknown mechanism driving *DLK1* expression through a super enhancer in a subset of neuroblastomas associated with high *DLK1* expression at both the RNA and protein levels. In addition, we show binding of CRC transcription factors associated with the adrenergic cell state up and downstream of the *DLK1* gene locus. Although chromatin interaction data would be required to show direct interaction with the *DLK1* promoter, it has been documented that DLK1 is associated with the adrenergic cell state in neuroblastoma ^54–58,72^. Orthogonal approaches were used to validate DLK1 expression on the surface of neuroblastoma and preferential expression compared to normal tissues. shRNA-based silencing and ATRA studies demonstrated a role of DLK1 in cellular differentiation, suggesting DLK1 influences the pluripotent state and differentiation block essential for cancer cell growth. *DLK1* mRNA expression was examined across cancer types where multiple subsets of samples with high expression was observed. Recent studies have reported DLK1 protein expression in several malignancies including endocrine/neuroendocrine, gastrointestinal, liver, lung, and renal tumors ^71^. We speculate the presence of a super enhancer may drive expression across multiple cancer histologies.

Here, we show that ADCT-701, a PBD dimer-based ADC targeting DLK1, was potently efficacious across all DLK1-expressing neuroblastoma xenograft models. IHC staining of DLK1 in our xenograft tumors highlights cellular heterogeneity ranging from high homogenous expression, more intermediate heterogenous patterns, and no protein present. Models exhibited a DLK1 expression-dependent response to ADCT-701, with high DLK1-expressing models having a complete or maintained complete response while no or low DLK1-expressing models enduring progressive disease. These data were utilized to support development of a Phase 1 first-in-human DLK1-directed immunotherapy clinical trial using ADCT-701 to treat neuroendocrine neoplasms, including neuroblastoma, in adult patients (ClinicalTrials.gov: NCT06041516). Neuroblastoma tumors are thought to include a mixture of adrenergic and mesenchymal cell states, with DLK1 likely being expressed predominately in the adrenergic subset. Since PBD dimers can penetrate surrounding cells, they can impact neighboring mesenchymal cells and those that do not express DLK1. ADCT-701 can have a positive clinical impact on heterogenous tumors; however, DLK1 negative relapses were observed in some cases. In addition, it is unknown if an antibody that binds will be internalized in a differentiated mature cell at a different rate than a proliferating cancer cell. Advances in bispecific immunotherapeutic approaches incorporating Boolean logic gates offer the potential for effectively targeting DLK1 on cancer cells while ameliorating potential toxicity due to adrenal and pituitary gland expression ^73^. This approach can build on a recent study focused on DLK1-directed CAR T cells if toxicity is observed ^74^.

Taken together, this study demonstrates the power of an integrative multi-omic approach to identify new immunotherapeutic targets and provides important validation and pre-clinical data to support a first-in-human DLK1-directed immunotherapy clinical trial for adult patients diagnosed with neuroendocrine neoplasms, including neuroblastoma (ClinicalTrials.gov: NCT06041516). Application of this approach to additional high-risk malignancies may reveal other surface proteins that can be targeted across cancers or cancer subtypes.

## STAR Table

| Antibodies | Source | Identifier |
| --- | --- | --- |
| DLK1-IHC | Abcam/ Santa Cruz | ab21682/ B7 |
| DLK1-Flow Cytometry | AdipoGen | AG-20A-0070-C100 |
| DLK1-Western | Cell Signaling | 2069S |
| KU80 | Cell Signaling | 2053S |
| Cadherin | Abcam | Ab6529 |
| Secondary IF Alexa | Thermo Fisher Scientific | Alexa Fluor 488 (A21206); Alexa Fluor 594 (A21203) |
| Goat anti-Mouse IgG (H+L) Cross-Adsorbed Secondary Antibody, PE | Thermo Fisher Scientific | P852 |
| ADCT-701 Antibody | ADCT | - |
| Reagent or Resource | Source | Identifier |
| Polybrene | Santa Cruz | SC-134220 |
| Puromycin | Santa Cruz | SC-108071 |
| ATRA | Sigma Aldrich | R2625-100mg |
| DAPI | Thermo Fisher Scientific | D3561 |
| Biological Samples | Source | Identifier |
| Human Neuroblastoma Primary Tumor Microarray (TMA) | Children's Hospital of Philadelphia (CHOP) | - |
| Human Neuroblastoma Patient Derived Xenograft Microarray (TMA) | Children's Hospital of Philadelphia (CHOP) | - |
| Pediatric Normal Tissue Microarray (TMA) | Children's Hospital of Philadelphia (CHOP) | - |
| Brain Tumor Mixed (TMA) | University of British Columbia (UBC) | - |
| Medulloblastoma (TMA) | University of British Columbia (UBC) | - |
| Mixed Tumor (TMA) | University of British Columbia (UBC) | - |
| Sarcoma Mixed (TMA) | University of British Columbia (UBC) | - |
| Sarcoma Mixed II (TMA) | University of British Columbia (UBC) | - |
| Critical Commercial Assay | Source | Identifier |
| SuperScript™ First Strand Synthesis for RT-PCR | Thermo Fisher Scientific | 11904018 |
| RNeasy Mini Kit | Qiagen | 74104 |
| QIAshredder | Qiagen | 79656 |
| HighSpeed Midi Kit | Qiagen | 12643 |
| FuGene 6 | Promega | E2691 |
| Lipofectamine | Thermo Fisher Scientific | L3000-008 |
| Experimental Models: Cell Lines | Source | Identifier |
| SK-N-Be(2)C | CHOP Cell Line Bank | - |
| Kelly | CHOP Cell Line Bank | - |
| NB-69 | CHOP Cell Line Bank | - |
| NLF | CHOP Cell Line Bank | - |
| SH-SY5Y | CHOP Cell Line Bank | - |
| Oligonucleotides | Source | Identifier |
| DLK1 MISSION shRNA Plasmid | Sigma Aldrich | TRCN0000055978 |
| DLK1 MISSION shRNA Plasmid | Sigma Aldrich | TRCN0000055979 |
| HPRT1 TaqMan Gene Expression Assay | Thermo Fisher Scientific | 4331182 - Hs02800695_m1 |
| GAPDH TaqMan Gene Expression Assay | Thermo Fisher Scientific | 4331182-Hs02758991_g1 |
| DLK1 TaqMan Gene Expression Assay | Thermo Fisher Scientific | 4331182-Hs00171584_m1 |
| Recombinant DNA | Source | Identifier |
| pMD2.G | Addgene Plasmid 11259 | https://www.addgene.org/12259/ |
| psPAX2 | Addgene Plasmid 11260 | https://www.addgene.org/12260/ |
| Software and Algorithms | Source | Identifier |
| STAR | Dobin et al 2013 | https://github.com/alexdobin/STAR |
| RSEM | Li and Dewey, 2011 | https://github.com/deweylab/RSEM |
| ROSE | Warren et al 2013; Loven et al 2013 | https://bitbucket.org/young_computation/rose/src/master/ |
| LILY | Boeva et al, in prep | https://github.com/BoevaLab/LILY |
| MaxQuant | Tyanova et al 2016 | https://www.maxquant.org/maxquant/ |
| Compartments | http://compartments.jensenlab.org/search | - |
| FlowJo | FlowJo LLC | http://www.flowjo.com |

## STAR Methods

### Neuroblastoma cell culture

Neuroblastoma cell lines (**Table 1**) were short tandem repeat typed (AmpFLSTR Identifier Kit – Thermo Scientific) and mycoplasma tested prior to use. Cells were grown in RPMI supplemented with 10% FBS and1% L-Glutamine at 37C and 5% CO_2_ (RPMI Complete). For plasma membrane protein isolation, cells were grown on ten 15cm plates per replicate and this was performed in duplicate. When cells reached 90% confluence, media was removed, and cells were washed with PBS. One plate was detached into single-cell suspension and counted to estimate the number of plates needed to reach 100M cells. Once PBS was removed and 3 ml of media were added, cells were scraped from the plate, collected in a 50mL conical and spun at 300 RCF for five minutes. Cell pellet was washed twice with PBS, supernatant was removed, and pellets were stored at −80 freezer.

### Neuroblastoma xenograft models

Neuroblastoma xenograft models (**Table 1**) were grown in CB17SC scid −/− female (5-8 weeks) mice following protocols established by the Institutional Animal Care and Use Committee (IACUC) at the Children’s Hospital of Philadelphia. Tumors were then removed from mice, dissected into 250 mg sections, washed with PBS, snap frozen in liquid nitrogen, and stored at −80 freezer. ^23^. More details on xenograft models are described in Rokita et al 2019 ^49^.

### Plasma membrane protein isolation

Plasma membrane surface protein isolation was modified from a prior study ^39^. Briefly, cells were resuspended in Homogenization Buffer (For 1000mL – 85.58g sucrose, 2.38g HEPES, 2mL 0.5M EDTA, pH 7.4 and Filter solution through 0.45 um polyethersulfone membrane bottle top filter) and passed through an 18-gauge needle three times followed by a 22-gauge needle three times. Next, cell lysis was processed with a Dounce homogenizer 40 times or dissociated using TissueRuptor (Qiagen). The sample was transferred to a 15mL conical and centrifuged at 1,000g for 10 minutes at 4C. Supernatant was loaded atop prepared ultracentrifugation tubes (13.2mL Beckman Cat#: BK344059) containing 2mL 60% sucrose. Homogenization buffer was used to balance the tubes for centrifugation at 100,000g for 60 minutes at 4C. Membrane proteins appeared as a white layer above the 60% sucrose, were collected and placed in a new ultracentrifugation tube. Six layers of sucrose (1.5mL each) were added in the following order: 42.8%, 42.3%, 41.8%, 41%, 39% and 37%. The tubes were balanced with 37% sucrose and subjected to ultracentrifugation at 100,000g for 18 hours at 4 degrees. Plasma membrane proteins were extracted from the top layer and placed in a new centrifuge tube. Each tube was filled with HEPES buffer (115mM NaCl, 1.2mM CaCl_2_, 1.2mM MgCl_2_, 2.4mM K_2_HPO_4_ and 20mM HEPES; pH 7.4) and samples were run at 150,000g to pellet the plasma membrane proteins. Tubes were decanted and proteins were placed in digestion buffer consisting of 1% sodium deoxycholate (Sigma Cat#: D6750-10G) in 100mM ammonium bicarbonate. Plasma membrane proteins were diluted 1:4 with 50mM ammonium bicarbonate and passed through 10K spin filters (Millipore Cat#: UFC501096). Samples were washed with water and protein was reduced in 5mM DTT for 1 hour at room temperature. Next, proteins were alkylated with 20mM iodoacetamide in the dark for 30 minutes and then digested with Trypsin (Promega, Cat V5113) overnight at room temperature. Samples were acidified using trifluoroacetic acid (TFA) at room temperature for 30 minutes and centrifuged at 14,000g for 20 minutes at 4 degree. Supernatant was desalted using C18 (Fisher Scientific, Cat 13-110-019) and eluted sample was dried using a speedvac.

### Mass spectrometry data generation

Dried samples were resuspended in 0.1% formic acid or 0.1% TFA if a TRAP column was utilized for mass spectrometry analysis. Cell line and xenograft plasma membrane proteins were analyzed using the Easy-nLC (Thermo Scientific) and Orbitrap Fusion Tribrid mass spectrometer (Thermo Scientific, San Jose CA). Peptides were separated using a column (75uM inner diameter) packed in the laboratory with C18 resin. Cell line and xenograft samples were analyzed using 135- and 165-minute gradient, respectively. The gradient ranged from 3% to 38% for 120 minutes; 38% to 98% for fifteen minutes (80% acetonitrile/0.1% formic acid) at a flow rate of 400-500nL/minute. The data was recorded using data-dependent acquisition (DDA). For full MS scan, the mass range of 350–1200 m/z was analyzed in the Orbitrap at 120,000FWHM (200 m/z) resolution and 5×10e5 AGC target value. High Collision Dissociation (HCD) collision energy was set to 32, AGC target to1×10e4 and maximum injection time to 120 msec. Detection of MS/MS fragment ions was performed in the ion trap.

### Mass spectrometry data analysis

Mass spectrometry raw files were searched using MaxQuant version 1.6.0.0 ^75^. MS/MS were searched with Andromeda using the Human UniProt FASTA database (July 2019). During the search, variable modifications were specified as methionine oxidation and N-terminal acetylation while fixed modification included carbamidomethyl cysteine. Trypsin, which cleaves after Lysine (R) and Arginine (K) was indicated as the digestive enzyme, with two permitted miscleavages. One or more razor or unique peptides were needed for protein identification and intensity based absolute quantification (iBAQ) was utilized for label-free quantification. False discovery rate was set at the peptide level (FDR < 0.01) and all other settings were standard. To account for injection amount, iBAQ values were log_2_ transformed after data was shown to be normally distributed and then further normalized to the mean across xenografts and cell lines. Missing values were imputed using Perseus ^76,77^ and used for all downstream analyses except surfaceome reproducibility and protein-RNA correlation.

### Multi-omic data analysis from cancer and normal tissues

Neuroblastoma cell line and xenografts RNA-sequencing (RNA-Seq) data were analyzed as described previously [46, 70]. Neuroblastoma patient and normal tissue (GTEx) RNA-Seq data were obtained from the Open Pediatric Cancer Project (OpenPedCan) (https://zenodo.org/record/7893332; https://d3b-center.github.io/OpenPedCan-methods/v/bbbb01b204201588f89d655a107f66b798a2ad18/). Cell line and xenograft models were processed with the same pipeline. This data allowed target expression and *MYCN* amplification status to be determined. ChIP-sequencing data was obtained from Upton et al 2020. Enhancer calls were made using LILY algorithms that builds on ROSE with correction for copy number ^59,78,79^. Epigenetic data were manually reviewed at the *DLK1* locus, resulting in assignment of a *DLK1* super-enhancer in SK-N-Be(2)C; this super-enhancer was initially annotated to a nearby gene using the LILY. Copy number data was calculated from svpluscnv ^80^. we utilized RNA-sequencing data from The Cancer Genome Atlas (TCGA) and Treehouse Childhood Cancer Initiative (https://treehousegenomics.soe.ucsc.edu/public-data/) to examine expression across cancer types.

### Data integration analysis

Mass spectrometry data was evaluated following log_2_ mean centering normalization. Next, the mean for each model was calculated and missing values were imputed using Perseus ^76,77^ for all analysis except reproducibility and correlation. In parallel, RNA data that was obtained from Open Targets was analyzed by calculating the mean for each tissue/histotype transcriptome wide. MS and RNA data were filtered using the annotations obtained from COMPARTMENTS (All Channel Integrated October 2020; GO Terms: GO:0005886|GO:0009279|GO:0009986|GO:0009897|GO:0005576; Confidence > 3) and surface protein consensus (SPC) score (December 2020; Score >=1) ^40,41,81^. Next, we evaluated normal tissue expression by calculating the fold change between neuroblastoma and all normal tissues except reproductive organs/brain and then we used a fold change cutoff of 4 as a read out of expression. We prioritized proteins that were present in less than 4 tissues and cell lines and in the top 50% abundance of either cell lines or xenografts. These proteins were visualized using an annotated heatmap that was designed using Complex Heatmaps ^82^.

### RNA-protein correlation analysis

Pearson correlation between RNA and protein was calculated following filtering with COMPARTMENTS and SPC as outlined above. Proteins that were only detected in two cells lines or xenografts were removed before generating the correlation value histogram. In parallel, a gaussian mixture model was used to identify two normal distributions in the RNA data. Those that had values below the mean of the lower distribution were extracted from the mass spectrometry data. Recurrent proteins are visualized using a bar plot.

### *MYCN* and *DLK1* correlation analysis

We investigated a role of *MYCN* in *DLK1* expression in neuroblastoma dataset from Therapeutically Applicable Research to Generate Effective Treatments (TARGET), SEQC and Kocak. Cohorts were analyzed in R2: Genomics Analysis and Visualization Platform (R2; http://r2.amc.nl) after selecting either Children’s Oncology Group (COG) high-risk or International Neuroblastoma Staging System (INSS) stage 4 patients (Kocak: n = 148; SEQC: n = 176; TARGET: n = 217).

### Overall survival analysis

A Kaplan Meier scan for overall survival was performed using R2: Genomics Analysis and Visualization Platform (R2; http://r2.amc.nl) using the median of *DLK1* expression to define groups for comparison. Prior to analysis, patients were first selected to include only high-risk or INSS stage 4 patients.

### Western blotting

Cells were detached with versine, collected, and washed using ice cold PBS and centrifuged at 300g for five minutes. Supernatant was removed and cells were resuspended in lysis buffer (25mM Tris Base, 150nM NaCl, 1mM EGTA, 1mM EDTA, 10mM NaF, 1% TritonX-100, 1mM DTT and Protease/Phosphatase Inhibitors (Cell Signaling #5872). Cells were lysed on ice for 30 minutes and sonicated briefly to aid lysis before centrifugation (18,000g at 4C for 15 minutes). Supernatant was reserved, aliquoted and stored at −80 for further use. Protein was quantified using a BCA assay (Pierce) and 30ug was run on a 8-16% Tris-Glycine gel, transferred to PVDF at 30V for 2 hours at room temperature or 15V overnight at 4C. Membrane was blocked for one hour in 5% milk in TBST and followed by overnight incubation with primary. Next, the membrane was washed three times with TBST for ten minutes, incubated with secondary for one hour and washed again three times with TBST. Blots were developed using SuperSignal West Femto Substrate (Cat #PI34095). Antibodies used in the study include: DLK1 (Cell Signaling – 2069S; 1:1000) and Ku80 (Cell Signaling – 2753S; 1:2500).

### Immunofluorescence

A total of 16,000 Kelly, SK-N-Be(2)C, SH-SY5Y, NB-69 and NLF cells were plated on 8 well chamber slides (NUNC, cat#154534). 48 hours later, cells were washed with PBS and fixed using 4% Formaldehyde (Pierce) for 10 minutes at room temperature. Next, cells were washed with ice cold PBS and blocked in PBST with 1% BSA for 30 minutes. Primary antibodies (DLK1-1:250; Cadherin-1:500) were incubated overnight at 4C and the next morning cells were washed twice with PBS for 10 minutes. Alexafluor secondary antibodies were applied for one hour in the dark and then the slide was washed twice with PBS for 10 minutes. DAPI was used to stain nuclei (1:5000 or 1ug/mL) at room temperature for 15 minutes and washed twice with PBS for 10 minutes. Slides were mounted using Prolong Diamond Mounting media and visualized on a Leica DM5000 B microscope with DFC365 FX camera.

### Immunohistochemistry

#### CHOP

DLK1 (Abcam, ab21682, Santa Cruz, B-7) antibody was used to stain formalin fixed paraffin embedded tissue slides. Staining was performed on a Bond Rx automated staining system (Leica Biosystems). The Bond Refine polymer staining kit (Leica Biosystems DS9800) was used. The standard protocol was followed with the exception of the primary antibody incubation which was extended to 1hr at room temperature and the post primary step was excluded. DLK1 was used at 1:2K dilution and antigen retrieval was performed with E1 (Leica Biosystems) retrieval solution for 20min. Slides were rinsed, dehydrated thru a series of ascending concentrations of ethanol and xylene, then coverslipped. Stained slides were then digitally scanned at 20x magnification on an Aperio CS-O slide scanner (Leica Biosystems).

#### UBC

Formalin-fixed, paraffin-embedded TMA sections were analyzed for DLK1 expression. In brief, tissue sections were incubated in Tris EDTA buffer (cell conditioning 1; CC1 standard) at 95C for 1 hour to retrieve antigenicity, followed by incubation with DLK1 antibody (Abcam ab21682) at 1:10,000 for 1 hour. Slides were then incubated with the respective secondary antibody (Jackson Laboratories) with 1:500 dilution, followed by Ultramap HRP and Chromomap DAB detection. For staining optimization and to control for staining specificity, normal organs were used as negative and positive controls. Intensity scoring was done on a common four-point scale. Descriptively, 0 represents no staining, 1 represents low but detectable degree of staining, 2 represents clearly positive staining, and 3 represents strong expression. Expression was quantified as H-Score, the product of staining intensity, and % of stained cells.

### Flow cytometry

Cells were collected and counted using the Cellometer Auto T4 (Nexcelom Bioscience) and one million cells per condition were placed in a FACS tube. Cells were centrifuged at 300g at 4C for five minutes and washed with PBS. To determine cell viability, Violet Viability Dye (Thermo Fisher, Cat# L34963) was applied to cells for 30 minutes in the dark. Cells were pelleted, washed with PBS and aliquot to either unstained, secondary only or DLK1. DLK1 cells were resuspended in DLK1 primary antibody (AG-20A-0070-C100 AdipoGen; 1:100 per million cells) for 30 minutes in the dark. After incubation, samples were washed twice with 1mL of FACS Buffer (PBS, 2% FBS, 1% EDTA) and then cells from DLK1 and secondary only were incubated with secondary antibody (P852; 1:500) protected from light for 30 minutes. Samples were washed twice with 1mL of FACS Buffer, then fixed in 200uL of 1-4% formaldehyde and stored at 4C in the dark until they were analyzed using a CytoFlex S (Beckman Coulter). Data was analyzed with FlowJo (BD Medical Technology).

### LentiVirus infection

Plasmid were obtained from Sigma as glycerol stocks (shDLK1: TRCN0000055978 and TRCN0000055979). Plasmid DNA was transfected using either Lipofectamine 3000 or Fugene into HEK293T cells. One day following transfection, media was changed to destination media (RPMI Complete). Infectious viral media was collected every 24 hours for two days and filtered through 0.45μm PVDF (Millipore Cat#: SE1M003M00). Viral media was combined with polybrene (Santa Cruz, Cat#: SC-134220) at 8 μg/mL media to increase transduction efficiency and applied to cells. 24 hours after infection, media on transduced cells was changed to RPMI complete. The next day, cells were selected with puromycin at the following concentrations: SK-N-Be(2)C (2400ng). When the mock plate completed selection, cells were harvested and knockdown was assessed by Western blotting.

### Cell proliferation and neurite analysis

Transduced cells were plated in a 96-well plate (Greiner-655098) at optimized plating densities (SK-N-Be(2)C-4,000 or 8,000/well; SH-SY5Y-12,000/well) and growth monitored in the Incucyte Zoom (Essen BioScience) with readings taken every hour. Six technical replicates were performed for each condition. Cell confluence and neurite outgrowth were quantified using optimized definitions for each cell type. P-values were calculated using a t-test, and representative images were selected for display purposes.

### All-trans retinoic acid (ATRA) treatment

Cells were plated in 6 well format and allowed to settle for 24 hours. Media was changed to 3% serum media for three hours. Media was then changed to either 3% serum or 3% serum supplemented with 5uM DMSO or ATRA. Media was changed every 48 hours until harvesting. SK-N-Be(2)C cells were harvested on day 5 and SH-SY5Y were harvested on day 7 post plating. Westerns were performed as outlined above. Cells were plated in 96 well format and treated in parallel. Representative images were extracted from the incucyte at the same time point as the cells collected for protein.

### Full proteome data generation and analysis

Cell pellets (obtained from a 10cm plate) were resuspended in 100uL of 6M Urea and 2M Thiourea (Sigma) with protease and phosphatase inhibitors (Cell Signaling 5872S). Samples were reduced in 5mM DTT for 1 hour at room temperature. Next, proteins were alkylated with 20mM iodoacetamide (IAA) in the dark for 30 minutes and then digested with Trypsin (Promega Cat#: PRV113) overnight at room temperature (1ug Trypsin: 20ug protein determined by BCA (Pierce Cat# 23227). Samples were dried and resuspended in 1% TFA and pH was adjusted if necessary to 2.0. Samples were desalted using C18 and eluted sample was dried using a speedvac. Samples were resuspended and processed by MS methods outlined above. Scrambled control and shDLK1 were performed in duplicate and injected on the MS twice. Raw files were database searched using MaxQuant as outlined in mass spectrometry data analysis section. T-test was performed using T.Test R package to assess differential expression. Proteins that were statistically significant (P < 0.05) and has a fold change of <−1 or > 1 were uploaded to Ingenuity pathway analysis.

### PK Studies

Quantification of total antibody and of conjugated antibody (drug-to-antibody ratio [DAR]>1) were determined using optimized electrochemiluminescence immunoassays (ECLIA). Calibration curves, quality controls, and study samples were diluted and added to either a soluble recombinant human DLK1 directly coated to an Meso Scale discovery (MSD) plate or to a biotinylated anti-PBD antibody coated to a streptavidin MSD plate respectively. The plates were then incubated and washed. A Sulfo-tag–labeled anti-Fc detector antibody (Sanquin) was used for measuring the total soluble DLK1 binding antibody. For the measuring of PBD-conjugated antibody a Sulfo-tag-labeled anti-HuBA-1-3D detector antibody (ADC Therapeutics) was used. For both assays after read buffer (MSD) was added, and the plates were read on the MSD QuickPlex Plate Reader (6000 Sector Imager; Meso Scale discoveryMSD).

### ADCT-701 mouse studies

CB17 SCID female mice were engrafted with tumors between five and seven weeks of age. Mice were randomized to one of three intervention arms approximately 2.5 weeks later when tumors reached 0.2 cm^3^ to: 1) 1 mg/kg ADCT-701; 2) 1 mg/kg B12-PL1601 (isotype, non-targeting PBD-conjugated ADC with same warhead); or 3) x mL phosphate buffered saline. If tumors on the ADCT-701 responded to treatment and regrew a second dose was administered. Mice were followed for up to 100 days or euthanized when tumor volumes reached of 2.0cm^3^. Thirteen CDX/PDX models with varying DLK1 expression were treated and evaluated. Study outcomes were determined using the criteria outlined by the NCI Pediatric Preclinical in Vivo Testing Program (PIVOT). Progressive disease is defined as either <50% tumor regression throughout study and >25% tumor growth by end of study (PD), mouse’s time to event <= 200% the Kaplan Meier (KM) median time-to-event in the control group (PD1) or mouse’s time to event >200% the KM median time-to-event in the control group (PD2). Stable Disease (SD) is determined when <50% tumor regression throughout the study but <25% tumor growth by end of the study. Partial response (PR) is recorded when >=50% tumor regression at some point during the study but measurable throughout the study. Complete Response (CR) is disappearance of measurable tumor mass during the study period. Lastly, a maintained complete response (MCR) is defined as no measurable mass for at least three consecutive weekly readings at any time after the treatment has been completed.

## Acknowledgements

This work was supported by a grant from the W.W. Smith Charitable Trust (SJD), an Innovation Award from Alex’s Lemonade Stand Foundation (SJD), and a Stand Up 2 Cancer-St. Baldrick¹s Pediatric Dream Team Translational Research Grant SU2C-AACR-DT1113 (JMM). This work was also supported by National Institutes of Health (NIH) grants U54-CA232568 (JMM), R01-CA204974 (SJD), R01-CA237562 (SJD), R03-CA230366 (SJD), U01-CA199287 (JMM), R35-CA220500 (JMM), U01-CA263957 (YPM), U01-CA199222 (MAS), F31-CA225069 (AKW), and T32-CA009140 (AKW). This work was delivered, in part, by the NexTGen Cancer Grand Challenges partnership funded by Cancer Research UK (CGCATF-2021/100002) and the National Cancer Institute (CA278687-01) and The Mark Foundation for Cancer Research.

## Author Contributions

Conceptualization, S.J.D., J.M.M; Methodology A.K.W., S.S., B.A.G., P.H.S. K.R.B., J.M.M, and S.J.D.; Validation, A.K.W., A.B.R., and K.L.C.; Formal Analysis, A.K.W., J.L.R.; J.P.,T.B., B.P., B.Z., C.Z., M.A.B., D.P.M; Investigation, A.K.W., A.B.R., M.T., D.M., K.L.C., M.V.L., Resources, B.T., S.W.E., F.Z., P.H.V., M.A.S., K.K., Y.P.M.; Data Curation, A.K.W., M.T., A.M., J.L.R., K.S.R., P.R.; Writing – Original Draft, A.K.W. and S.J.D.; Writing – Review and Editing, J.L.R, M.A.S., K.R.B., P.H.S., P.H.V, B.A.G., J.M.M, S.J.D.; Visualization, A.K.W. and A.B.R.; Project Administration, S.J.D. and J.M.M.; Supervision, B.A.G., J.M.M., S.J.D.; Funding Acquisition, A.K.W., B.A.G., J.M.M. and S.J.D.

## Declaration of Interests

F. Zammarchi, K. Havenith and P.H. van Berkel are or were employed by ADC Therapeutics at the time the work was conducted, and all hold or previously held shares/stocks in ADC Therapeutics

## Notes

### Summary of Updates

Fixed figures that exhibited low resolution; author affiliations updated; acknowledgements updated; contributions updated.

