## Supplementary Figures for "A proteogenomic surfaceome study identifies DLK1 as an immunotherapeutic target in neuroblastoma"

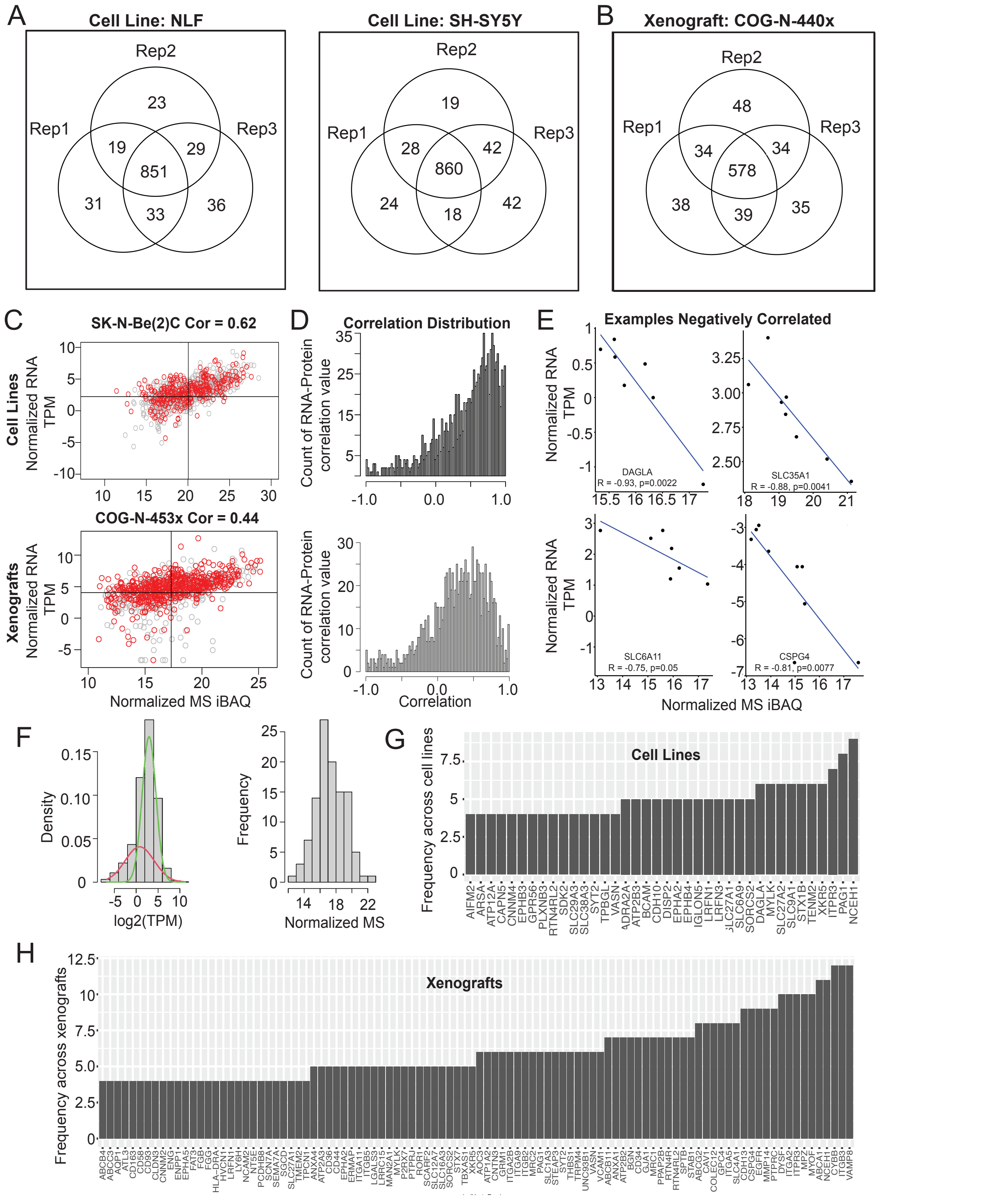


**Figure S1 Neuroblastoma cell line and xenograft model surfaceome analyses show strong correlation between replicates and confirm lack of correlation between RNA and protein; Related to Figure 1 (A)** Venn diagram of protein identified in three biological replicates (2 technical injections each) of NLF, SH-SY5Y. **(B)** Venn diagram of protein identified in three replicates of COG-N-440x. **(C)** Correlation between RNA and protein from the same cell line (SK-N-Be(2)C) and xenograft (COG-N-453x). **(D)** Global distribution of correlation values for cell lines and xenograft models. **(E)** Examples of negatively correlated RNA and proteins. **(F)** Gaussian mixture model (GMM) applied to the RNA-sequencing data and overlap of low RNA and detected by MS proteins. **(G-H)** Bar plots of recurrent low RNA and identified by mass spectrometry proteins for cell lines and xenografts.

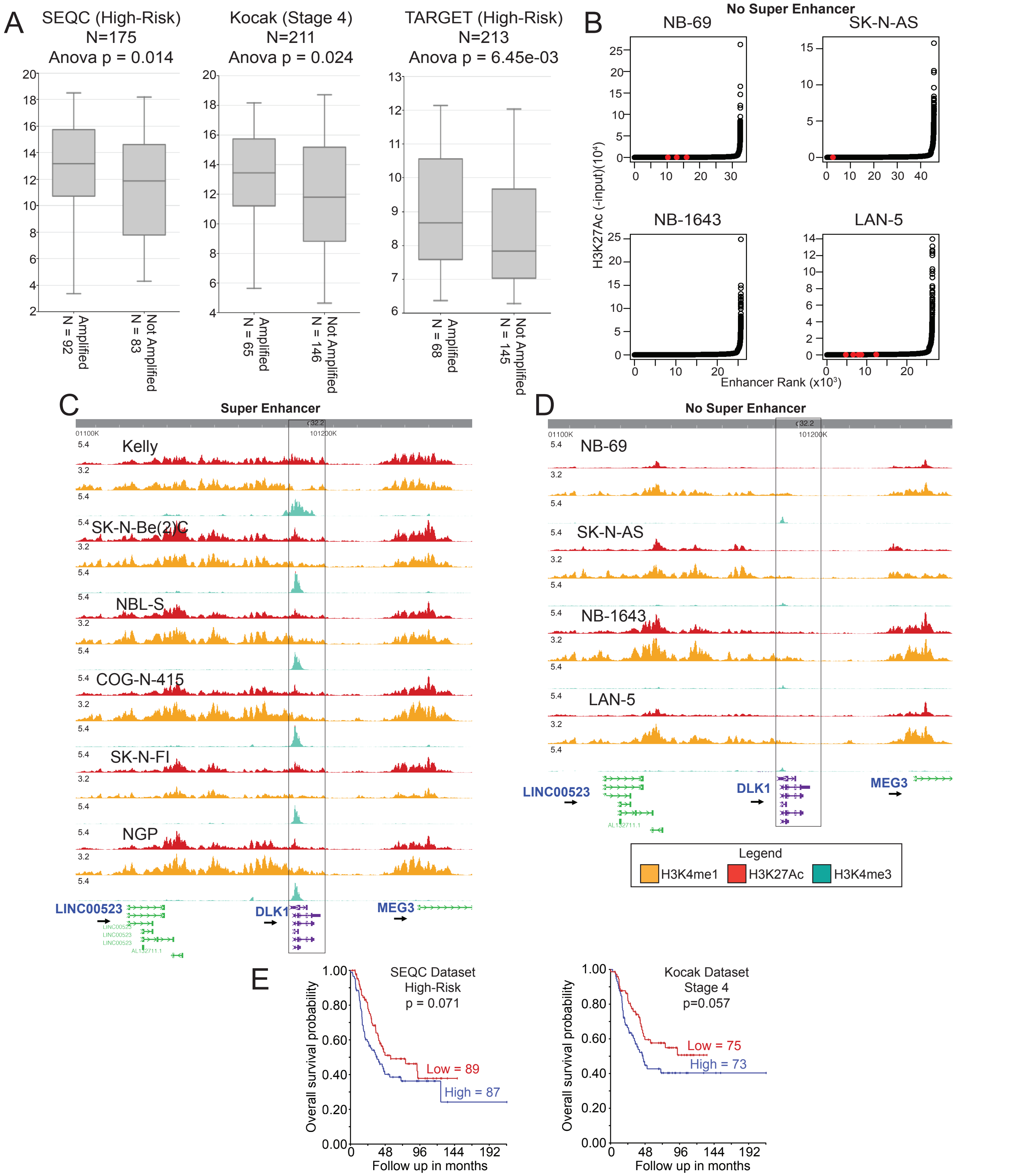


**Figure S2 DLK1 has a super enhancer, does not correlate with MYCN status, and correlates with poor outcome; Related to Figure 3 (A)** DLK1 and MYCN expression in high patients from three cohorts (TARGET High Risk, SEQC High Risk and Kocak Stage 4). **(B)** Swoosh plots ranking super enhancers by peak score percentage as called by LILY algorithm in a panel of cell lines. Cell lines shown do not have a super enhancer. **(C-D)** Visualization of H3K27Ac, H3K4me1 and H3K4me3 at the DLK1 locus in cell lines with and without the super enhancer in the WashU Epigenome Browser v52 1.0. **(E)** Kaplan Meier performed in R2 show high levels of DLK1 trend with worse outcome in two cohorts but did not reach statistical significance (SEQC and Kocak).

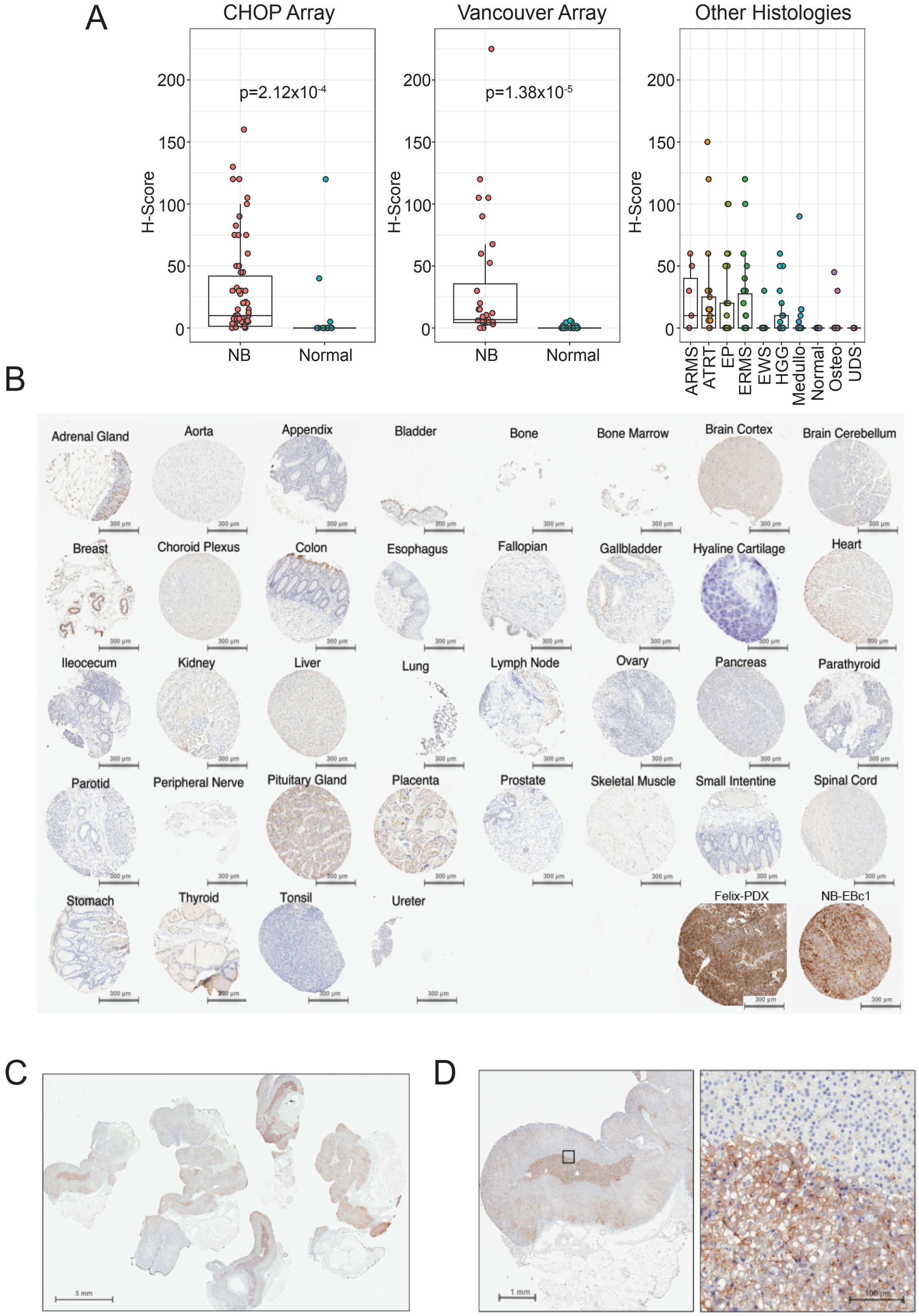


**Figure S3 DLK1 has limited expression in pediatric normal tissues as assessed by immunohistochemistry; Related to Figure 4 (A)** Differential expression of NB and normal tissues H-scores as scored from IHC microarrays from CHOP and Vancouver. Additional staining was performed in other histotypes **(B)** IHC of pediatric normal tissues to evaluate DLK1 expression. Two neuroblastoma tumors are displayed for comparison **(C)** Extensive staining of pediatric adrenal glands to assess DLK1 expression **(D)** IHC shows staining in the adrenal medulla compared to adrenal cortex. The left panel shows one adrenal gland displayed in Panel C and the right panel shows the region highlighted by the black box.

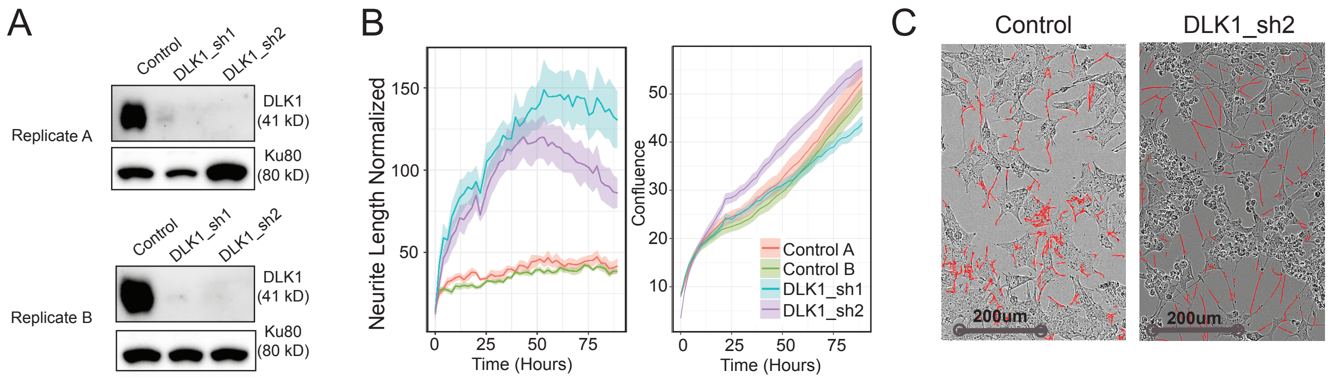


**Figure S4 DLK1 knockdown results in neurite formation; Related to Figure 5 (A)** Western blot of DLK1 and loading control Ku80 showing depletion of DLK1 in two replicates with two shRNAs. **(B)** Neurite outgrowth and confluence measured by Incucyte Zoom on control and knockdown cells using two shRNA. **(C)** Images of control and DLK1 knockdown cells.

A

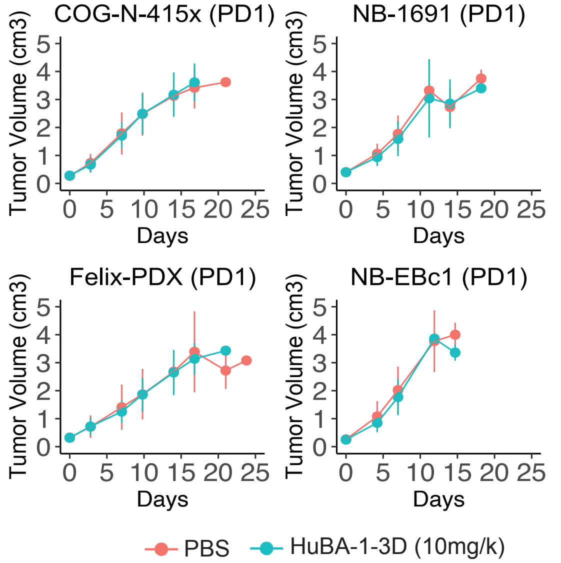


B

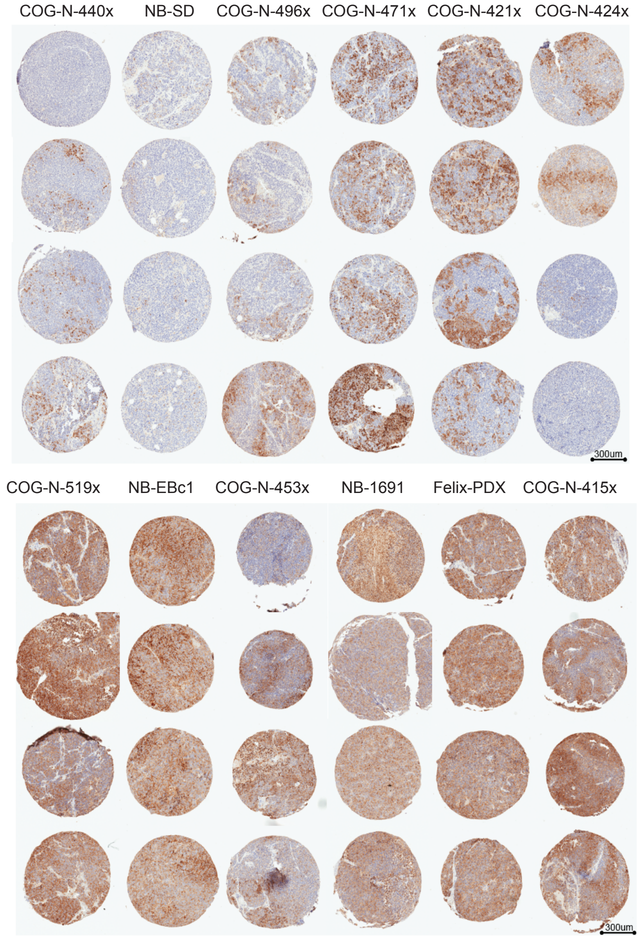


C

| Tumor | Grp | N | KM med | EFS T - C | EFS T/C | p-value Gehan-Wilcoxon | minRTV mean±SD | minRTV p-value | Med resp |
|---|---|---|---|---|---|---|---|---|---|
| COG-N-440x | Saline | 2 | 9.2 | | | | 1.348±0.560 | | PD |
| | ADCT-701 | 2 | 10.7 | 1.4 | 1.16 | p = 1.000 | 1.658±0.578 | p = 0.667 | PD1 |
| | B12-PL1601 | 2 | 7.2 | -2.1 | 0.77 | p = 1.000 | 1.916±0.272 | p = 0.667 | PD1 |
| COG-N-415x | Saline | 2 | 4.7 | | | | 2.887±0.648 | | PD |
| | ADCT-701 | 2 | > 101 | > 96.3 | > 21.32 | p = 0.102 | 0.000±0.000 | p = 0.333 | MCR |
| | B12-PL1601 | 2 | 5.8 | 1.1 | 1.22 | p = 0.450 | 2.312±0.219 | p = 0.667 | PD1 |
| COG-N-421x | Saline | 2 | 12.5 | | | | 1.098±0.340 | | PD |
| | ADCT-701 | 2 | 89.3 | 76.9 | 7.17 | p = 0.102 | 0.464±0.357 | p = 0.333 | SD |
| | B12-PL1601 | 2 | 20 | 7.6 | 1.61 | p = 0.102 | 0.813±0.048 | p = 0.333 | PD1 |
| COG-N-424x | Saline | 2 | 4.8 | | | | 2.612±0.625 | | PD |
| | ADCT-701 | 2 | > 102 | > 97.2 | > 21.18 | p = 0.102 | 0.000±0.000 | p = 0.333 | MCR |
| | C | 2 | 5.7 | 0.9 | 1.19 | p = 0.450 | 2.763±0.261 | p = 1.000 | PD1 |
| COG-N-452x | Saline | 2 | 7.9 | | | | 1.537±0.025 | | PD |
| | ADCT-701 | 2 | > 101 | > 93.1 | > 12.82 | p = 0.102 | 0.000±0.000 | p = 0.333 | MCR |
| | B12-PL1601 | 2 | 17.5 | 9.6 | 2.22 | p = 0.102 | 1.512±0.143 | p = 1.000 | PD1 |
| COG-N-453x | Saline | 2 | 8.9 | | | | 1.693±0.082 | | PD |
| | ADCT-701 | 2 | > 102 | > 93.1 | > 11.49 | p = 0.102 | 0.000±0.000 | p = 0.333 | MCR |
| | B12-PL1601 | 2 | 19.4 | 10.6 | 2.19 | p = 0.102 | 1.555±0.200 | p = 0.667 | PD1 |
| COG-N-471x | Saline | 2 | 10.4 | | | | 2.010±0.649 | | PD |
| | ADCT-701 | 2 | > 102 | > 91.6 | > 9.77 | p = 0.102 | 0.000±0.000 | p = 0.333 | CR |
| | B12-PL1601 | 2 | 13.4 | 2.9 | 1.28 | p = 0.102 | 1.797±0.538 | p = 0.667 | PD1 |
| COG-N-496x | Saline | 2 | 6.7 | | | | 2.121±0.230 | | PD |
| | ADCT-701 | 3 | 15 | 8.3 | 2.24 | p = 0.046 | 1.111±0.606 | p = 0.200 | PD2 |
| | B12-PL1601 | 2 | 5.7 | -1 | 0.86 | p = 0.102 | 2.427±0.756 | p = 1.000 | PD1 |
| COG-N-519x | Saline | 1 | 9.8 | | | | 0.995±NA | | PD |
| | ADCT-701 | 3 | 91.6 | 81.9 | 9.38 | p = 0.083 | 0.025±0.043 | p = 0.500 | MCR |
| | B12-PL1601 | 1 | 7.8 | -2 | 0.79 | p = 0.317 | 1.543±NA | p = 1.000 | PD1 |
| FelixPDX | Saline | 2 | 11.1 | | | | 1.335±0.219 | | PD |
| | ADCT-701 | 2 | 44.2 | 33.2 | 4 | p = 0.102 | 0.135±0.191 | p = 0.333 | CR |
| | B12-PL1601 | 2 | 9.8 | -1.3 | 0.89 | p = 0.102 | 1.664±0.024 | p = 0.333 | PD1 |
| NB-1691[c] | Saline | 2 | 5 | | | | 2.702±1.029 | | PD |
| | ADCT-701 | 2 | > 103 | > 98 | > 20.71 | p = 0.102 | 0.000±0.000 | p = 0.333 | CR |
| | B12-PL1601 | 2 | 4.7 | -0.3 | 0.94 | p = 1.000 | 2.864±0.938 | p = 0.667 | PD1 |
| NB-EBc1[c] | Saline | 2 | 8.3 | | | | 1.071±0.635 | | PD |
| | ADCT-701 | 2 | > 103 | > 94.7 | > 12.44 | p = 0.102 | 0.000±0.000 | p = 0.333 | MCR |
| | B12-PL1601 | 2 | 9.1 | 0.9 | 1.1 | p = 1.000 | 1.270±0.068 | p = 1.000 | PD1 |
| NB-SD[c] | Saline | 3 | 8 | | | | 1.803±0.172 | | PD |
| | ADCT-701 | 3 | 23 | 15 | 2.87 | p = 0.275 | 1.866±0.497 | p = 1.000 | PD2 |
| | B12-PL1601 | 3 | 10.6 | 2.6 | 1.33 | p = 0.275 | 1.308±0.348 | p = 0.200 | PD1 |

D

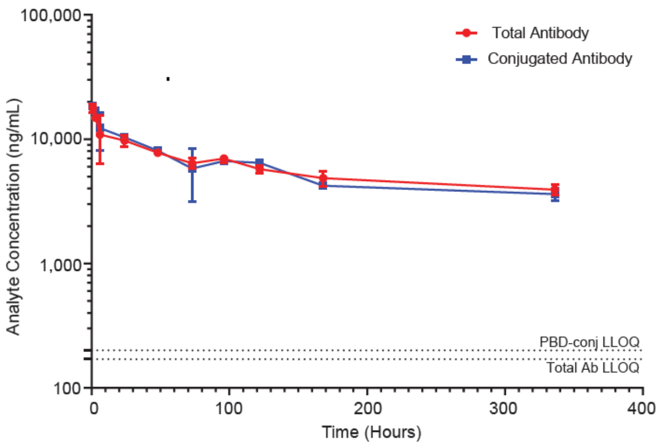


**Figure S5 ADCT-701 shows efficacy in xenograft models with high DLK1 expression which was not observed with the antibody lacking a PBD payload; Related to Figure 6:** (A) Preclinical in vivo studies with the antibody alone showed no efficacy in models with high expression of DLK1. (B) IHC staining on four cores per model for DLK1 efficacy studies. (C) Table of statistics related to mouse efficacy studies. (D) Serum concentrations (ng/mL; mean +/-SD) of total and conjugated antibody (DAR ≥1) in non-tumor-bearing mice following a single intravenous administration of ADCT-701 at 1 mg/kg. LLOQ (lower limit of quantification) is indicated for both total and conjugated antibody. KM med: Kaplan-Meier estimate of median time-to-event (days); EFS T-C: difference in median time-to-event (days) of treatment group compared to control group; EFS T/C: ratio of median time-to-event (days) of treatment group compared to control group; Min RTV p-value: computed using Wilcoxon rank sum test; Med resp: median response evaluation. PD1/PD2-Progressive Disease; SD-Stable Disease; CR-Complete Response; MCR-Maintained Complete Response.
